# ZeroCostDL4Mic: an open platform to use Deep-Learning in Microscopy

**DOI:** 10.1101/2020.03.20.000133

**Authors:** Lucas von Chamier, Romain F. Laine, Johanna Jukkala, Christoph Spahn, Daniel Krentzel, Elias Nehme, Martina Lerche, Sara Hernández-Pérez, Pieta K. Mattila, Eleni Karinou, Séamus Holden, Ahmet Can Solak, Alexander Krull, Tim-Oliver Buchholz, Martin L. Jones, Loïc A Royer, Christophe Leterrier, Yoav Shechtman, Florian Jug, Mike Heilemann, Guillaume Jacquemet, Ricardo Henriques

**Author notes:** Equal contributing authors (Lucas von Chamier, Romain F. Laine).

## Abstract

The resources and expertise needed to use Deep Learning (DL) in bioimaging remain significant barriers for most laboratories. We present https://github.com/HenriquesLab/ZeroCostDL4Mic/wiki, a platform simplifying access to DL by exploiting the free, cloud-based computational resources of Google Colab. https://github.com/HenriquesLab/ZeroCostDL4Mic/wiki allows researchers to train, evaluate, and apply key DL networks to perform tasks including segmentation, detection, denoising, restoration, resolution enhancement and image-to-image translation. We demonstrate the application of the platform to study multiple biological processes.

---

Despite the enthusiasm and innovations fuelled by DL technology, the need to access powerful and compatible resources, install multiple computational tools, and modify code to train neural networks all lead to a significant accessibility barrier that is difficult to cross by scientists without a strong background in computer science. This includes most biologists, clinicians, pathologists and microscopy users. To perform a specific bioimage analysis task, a typical DL pipeline requires users first to train a neural network with appropriate training data. Once trained, the DL network can be applied to analyze images that are similar to those used during training. Training is the crucial part of the DL pipeline as it will dictate the specificity and performance of the DL network (1, 2). However, it is also a challenging aspect of the process as it requires specialized knowledge and access to specialised computational resources. As a result, researchers may find it easier to use pre-trained networks available online to process their images. This approach may alleviate the onerous computational requirement of training and may even produce visually appealing imaging data. However, it has become clear that using pre-trained networks without appropriate additional training on the specific data of interest (a process known as transfer learning (3)), can lead to artifactual and misleading predictions (Supplementary Note 1).

Here, we present https://github.com/HenriquesLab/ZeroCostDL4Mic/wiki, an easy-to-use deployment DL platform which considerably simplifies the use of DL for microscopy (Supplementary Video 1). Importantly, ZeroCostDL4Mic allows researchers with little or no coding expertise to train (and re-train), validate and use DL networks (Figure 1). In parallel, it guides researchers to acquire more knowledge, to experiment with optimizing DL parameters and generate the training data necessary for DL. It uses Google Colab which provides the free, cloud-based computational resources needed for each step in the DL pipeline. We currently offer solutions to multiple powerful bioimage analysis tasks made possible with DL within Zero-CostDL4Mic (Figure 2 and Supplementary Video 2). These include image segmentation and object detection (using U-Net (4–6), StarDist (7, 8) and YOLOv2 (9)), image denoising and restoration (using CARE (10) and, Noise2Void (11)), super-resolution microscopy (using Deep-STORM (12)) and image-to-image translations (using Label-free prediction - fnet (13), pix2pix (14) and CycleGAN (15)) (see Supplementary Note 2, Supplementary Fig. 2-7 and Supplementary Video 2-11).

**Fig. 1.**
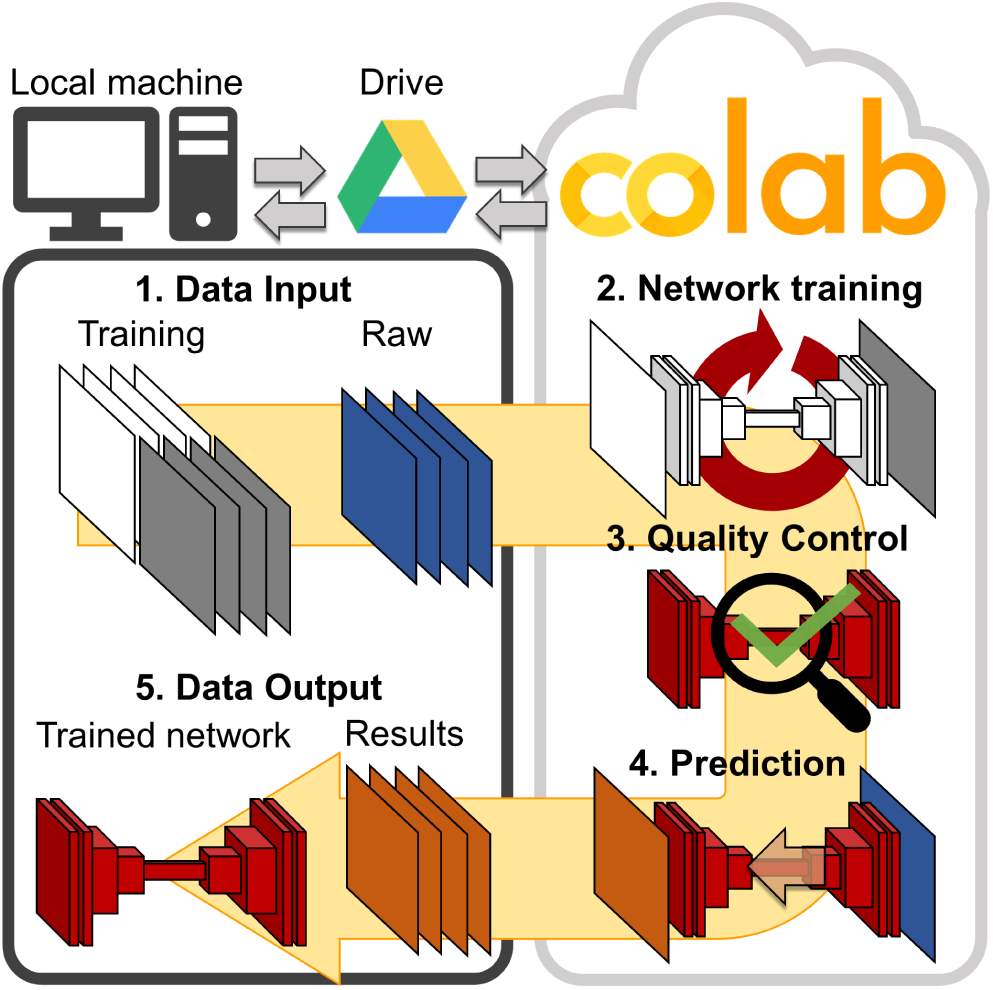
Overview of ZeroCostDL4Mic. Workflow of ZeroCostDL4Mic, featuring data transfer through Google Drive, plus training, quality control and prediction via Google Colab. After running a network, both trained models and prediction results can then be downloaded to the user’s machine.

**Fig. 2.**
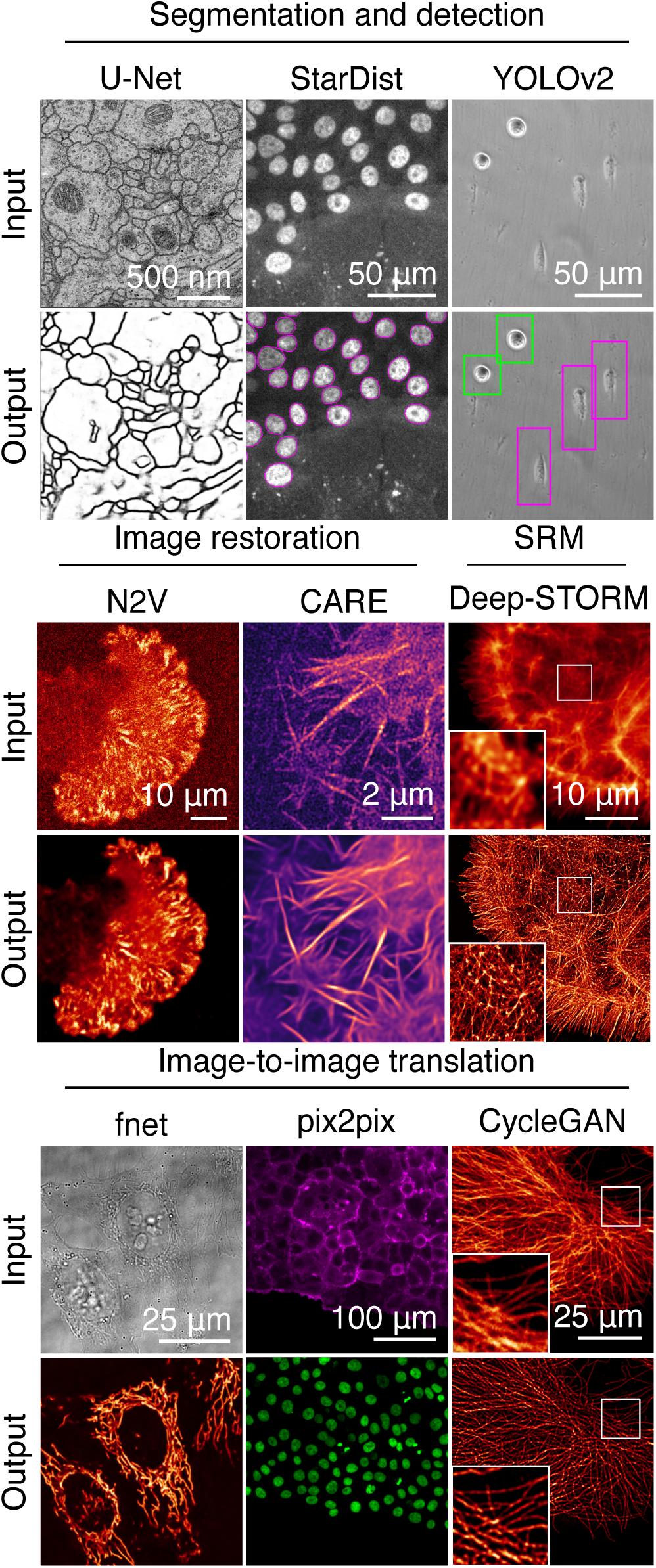
Overview of the bioimage analysis tasks currently implemented within ZeroCostDL4Mic platform. Datasets from top left to bottom right: U-Net – ISBI 2012 Neuronal Segmentation Dataset16, Stardist – nuclear marker (SiR-DNA) in DCIS.COM cells, YOLOv2 – brightfield in MDA-MB-231 cells, N2V – actin label (paxillin-GFP) in U-251-glioma cells, CARE – actin label Lifeact-RFP in DCIS.COM cells, Deep-STORM – actin-labeled glial cell, fnet – brightfield and mitochondrial label TOM20-AlexaFluor 594 in HeLa cells, pix2pix – actin label Lifeact-RFP and nuclear labels in DCIS.COM cells, CycleGAN – tubulin label in U2OS cells. All datasets are available through Zenodo or as indicated in the GitHub repository.

In practice, ZeroCostDL4Mic is a collection of self-explanatory Jupyter Notebooks, featuring an easy-to-use graphical user interface (GUI) (Supplementary Fig. 8) that requires only a web browser and a Google Drive account for a user to run any of our DL-based tasks. Jupyter Notebooks can efficiently and interactively run Python code, currently the default language to deploy DL applications. All calculations are performed in the cloud, circumventing the need to purchase or install graphical processing units (GPUs) and associated software.

Using ZeroCostDL4Mic does not require prior knowledge in coding. Researchers can, in a few mouse clicks and aided by a simple workflow, install all needed software dependencies, upload their imaging data and run networks for training and prediction (Supplementary Video 1). While the underlying code is hidden by default, it remains accessible, allowing users to learn, explore, and edit the notebooks’ programmatic structure.

ZeroCostDL4Mic notebooks share a common workflow to promote easy adoption and good practice in DL, encouraging users to use models trained on their data. Indeed, before deploying trained models on unseen data, users can easily test their quality (see Supplementary Note 3 and Supplementary Fig. 9 and 10). This is fundamental in optimizing the network performance for a particular application, determining its limitations and preventing the significant introduction of artifacts, a commonly raised concern for DL applications in microscopy (1, 2). In practice, we implemented a quantitative quality control step in all notebooks, which allows the assessment (discussed in Supplementary Note 3) and improvement of model performance (Supplementary Fig. 11).

Additionally, we enabled several important functionalities that facilitate and improve the applicability of our DL approaches. In particular, we implemented: (1) automated data augmentation which can artificially expand the image diversity of a dataset, especially beneficial when only small training datasets are available (Supplementary Note 4 and Supplementary Fig. 12 and 13); (2) transfer learning (3), allowing to take advantage of pre-trained networks (from so-called model “zoos”) by re-using previously learned features within these models and therefore speeding up and improving the training process (see Supplementary Note 5 and Supplementary Fig. 14 and 15); and (3) batch processing (predictions) of unseen data to streamline analysis once a satisfactory model has been obtained. It is important to note that trained models can also be downloaded and used outside ZeroCostDL4Mic (e.g., StarDist in Fiji(16)).

Via Google Colab, ZeroCostDL4Mic provides free access to the high-performance computational resources needed to run the large range of DL networks implemented here (Supplementary Note 2 for networks and Supplementary Note 6 for resources). For each featured network, we provide an example dataset that researchers can use to test and learn the basis of its workflow. For these datasets, we show that the corresponding training sessions only take a few minutes to a few hours (Supplementary Table 2), allowing the generation of high-performance DL models, producing the inference outputs shown in Figure 2 and the Supplementary Figures and Movies (Supplementary Fig. 2-6 and Supplementary Video 2-11. We also highlight the versatility and power of Zero-CostDL4Mic by sequentially combining multiple DL tasks such as image-to-image translation and tracking, and by integrating it within larger image analysis pipelines to enable, for instance, automated cell tracking(17) (Supplementary Fig. 7). Therefore, we envision ZeroCostDL4Mic constitutes an easily accessible and adaptable starting point to the use and deployment of DL-based bioimage analysis.

ZeroCostDL4Mic complements current community efforts to simplify access to DL in microscopy, e.g., ImJoy (18) and ilastik (19) or integration projects of DL into Fiji/ImageJ (7, 8, 10, 16, 20) it also substantially differs from these solutions by providing a single simple platform to carry out the necessary end-to-end DL workflow: install the various computational components, train a model using custom data, quantitatively validate the performance of the model and deployment on new data. By allowing training on custom data, we provide an alternative to the use of inappropriate pre-trained models, which often do not correctly represent the types of data researchers will want to analyze (Supplementary Note 1 and Supplementary Fig. 1).

By bringing previously published methods into a streamlined format that allows easy, cost-free access and customized use of DL in microscopy, we believe this resource is an important step towards widening the use of DL approaches beyond the community of computer scientists to the laboratories that generate the imaging data. In parallel, it enables researchers to improve their understanding of DL and experiment with optimizing DL parameters and choosing appropriate networks for a specific application. These steps are essential to both exploit the benefits and understand the limitations of DL approaches in research. We envision that the templates presented here can be used by DL developers to showcase their own network architectures in a unified and reproducible framework (Supplementary Note 7). This will ensure the rapid dissemination of novel technologies and provide consistent user experience for reproducible and comparative studies of DL approaches. Altogether, ZeroCostDL4Mic has the potential to dramatically accelerate the uptake of DL for new users and promotes their capacity to use increasingly sophisticated and powerful imaging analysis strategies.

## Availability

ZeroCostDL4Mic is available as Supplemental Software or can be accessed from our https://github.com/HenriquesLab/ZeroCostDL4Mic. This resource is fully open-source, providing users with tutorials, Jupyter Notebooks for Google Colab, and many real-life example datasets for training and testing. The example datasets are available for download in https://zenodo.org/search?page=1&size=20&q=ZeroCostDL4Mic (links provided in Supplementary Table 1 and our GitHub page).

## ACKNOWLEDGEMENTS

First and foremost, we would like to thank Dr. Martin Weigert from the Swiss Federal Institute of Technology (EPFL) and Dr. Uwe Schmidt from the Max Planck Institute of Molecular Cell Biology and Genetics (MPI-CBG), who pioneered a considerable portion of the technology this work is based on and whose ethos in making Deep Learning more accessible for microscopy helped inspire this work. Dr. Schmidt has kindly given key feedback during the preparation of the manuscript. All the network architectures and tasks presented here originate from already published work, having been edited and prepared for Google Colab to simplify their uptake by novice users. When using the ZeroCostDL4Mic platform, please cite the original publications associated with each network.

This work was funded by grants from the UK Medical Research Council (MR/K015826/1) (R.H.), the Wellcome Trust (203276/Z/16/Z) (R.H.) and the Gulbenkian Foundation (R.H.). R.F.L. would like to acknowledge the support of the MRC Skills development fellowship (MR/T027924/1). This work was also supported by grants awarded by the Academy of Finland (to G.J. and P.K.M.), the Sigrid Juselius Foundation (to G.J. and P.K.M.), the University of Turku foundation and Turku Doctoral Program in Molecular Medicine (TuDMM)(to SHP), Åbo Akademi University Research Foundation (CoE CellMech; G.J.) and by Drug Discovery and Diagnostics strategic funding to Åbo Akademi University (G.J.). We thank Dr. Aki Stubb for providing us with the raw data used to showcase Noise2Void 2D. The Cell Imaging and Cytometry Core facility (Turku Bioscience, University of Turku, Åbo Akademi University, and Biocenter Finland) and Turku Bioimaging are acknowledged for services and instrumentation and expertise. M.L is supported by Victoriastiftelsen (FI). A.K, T. - O.B and F.J. funded by German Research Foundation (DFG) under the code JU3110/1-1 and German Federal Ministry of Research and Education under the code 01IS18026C, ScaDS2. M.H. and C.S. acknowledge funding by the German Science Foundation (grant nr. SFB1177), C.S. additionally acknowledges support by the European Molecular Biology Organization (short term fellowship 8589). C.L. acknowledges the support of CNRS through the ATIP A0 2016 grant. SH and EK were supported by a Wellcome Trust Royal Society Sir Henry Dale Fellowship grant number 206670/Z/17/Z to SH. We additionally thank Chan Zuckerberg Biohub and its donors for funding Loic A. Royer and Ahmet Can Solak’s work. This work was supported by the Francis Crick Institute, which receives its core funding from Cancer Research UK (FC001999), the UK Medical Research Council (FC001999), and the Wellcome Trust (FC001999). Note: CoLaboratory™ and Google Drive™ are trademarks of Google LLC - ©2018 Google LLC All rights reserved.

## AUTHOR CONTRIBUTIONS

G.J. and R.H. conceived the project; L.v.C., R.F.L, J.J., D.K., E.N., T.-O.B. and G.J., and R.H. wrote source code based on the work of A.K., T.-O.B., F.J., E.N., Y.S., and L.A.R. among others; L.v.C., R.F.L., J.J., C.S., C.L., and G.J. performed the image acquisition of the test and example data; L.v.C., R.F.L., J.J., C.S., M.L., S.H.-P., P.K.M., E.K., S.H., A.K., T.-O. B., C.L., M.L.J., D.K., E.N., Y.S., M.H., and G.J. tested the platform; L.v.C., R.F.L., J.J., C.S., G.J. and R.H. wrote the manuscript with input from all co-authors.

