## Supplementary material for "ZeroCostDL4Mic: an open platform to use Deep-Learning in Microscopy": Online Methods

#### Cell Culture.

U-251 glioma cells were grown in DMEM/F-12 (Dulbecco's Modified Eagle's Medium/Nutrient Mixture F-12; Life Technologies, 10565-018) supplemented with 10 % fetal bovine serum (FCS) (Biowest, S1860). U-251 glioma cells expressing endogenously tagged paxillin-GFP were generated using CRISPR / Cas9 and were described previously[1].

MCF10 DCIS.COM (DCIS.COM) lifeact-RFP cells were cultured in a 1:1 mix of DMEM (Sigma-Aldrich) and F12 (Sigma-Aldrich) supplemented with 5% horse serum (16050-122; GIBCO BRL), 20 ng/ml human EGF (E9644; Sigma-Aldrich), 0.5 mg/ml hydrocortisone (H0888- 1G; Sigma-Aldrich), 100 ng/ml cholera toxin (C8052-1MG; Sigma-Aldrich), 10 µg/ml insulin (I9278-5ML; Sigma- Aldrich), and 1% (vol/vol) penicillin/streptomycin (P0781-100ML; Sigma-Aldrich). DCIS.COM lifeact-RFP cells were generated using lentiviruses, produced using pCDH-LifeAct mRFP, psPAX2, and pMD2.G constructs - see [2] for more details.

HeLa ATCC cells were seeded on fibronectin-coated 8-well chamber slides (Sarstedt, Germany, 1.5 x 104 cells/well). Cells were grown for 16h at 37°C and 5% $CO_2$ in Dulbecco's modified Eagle's medium containing 4.5 g/l glucose, 10% FCS, and 1% L-alanyl-L-glutamine (Thermo Fisher, GlutaMAX). To fix the HeLa cells, we employed a protocol shown to preserve the cytoskeleton and organelles (adapted from [3]). In brief, the culture medium was directly replaced with PHEM buffer containing 3% methanol-free formaldehyde (Thermo Fisher, USA) and 0.2% EM-grade glutaraldehyde (Electron Microscopy Sciences, USA) and incubated the samples for 1h at room temperature. Cells were washed thrice with PBS, quenched with 0.2% sodium borohydride in PBS for 7 min and washed again thrice with PBS.

A2780 cells were cultured in RPMI 1640 supplemented with 10% FCS. The cells were grown at 37°C in a 5% $CO_2$ incubator.

MDA-MB-231 (triple-negative human breast adenocarcinoma) cancer cells were grown in DMEM (Sigma-Aldrich) supplemented with 10% FCS at 37°C and 5% $CO_2$.

Rat hippocampal neurons from embryonic day 18 pups[4] were cultured on 18 mm coverslips at a density of 6,000 cells/cm$^2$ following established guidelines of the French Animal Care and Use Committee (French Law 2013-118 of 1st February 2013) and approval of the local ethics committee (agreement 2019041114431531-V2 #20242). In these neuronal cultures, a small number of glial cells, such as the one shown in Figure 2, Sup. Fig. 5 and Sup. Movie 10 are present and were labeled and imaged.

U2OS cells were purchased from DSMZ (Leibniz Institute DSMZ-German Collection of Microorganisms and Cell Cultures, Braunschweig DE, ACC 785). MDA-MB-231 cells were provided by ATCC. DCIS.COM were provided by J.F. Marshall (Barts Cancer Institute, Queen Mary University of London, London, England, UK). HeLa cells were provided by ECACC.

#### U-Net training dataset.

The training datasets used for segmentation in the Unet ZeroCostDL4Mic notebooks are publicly available EM datasets. For 2D segmentation, this was a neuronal membrane segmentation dataset from the ISBI challenge 2012[5] and for 3D segmentation from the mitochondrial segmentation dataset available from EPFL[6]. Datasets for segmentation tasks can also be created manually. This requires target images which have been segmented by an expert using drawing tools, e.g. in ImageJ/Fiji[7], to draw outlines around the structures of interest. For training in the notebook, the source (raw EM image) and target (8-bit mask obtained from expert drawing) images were placed in separate folders, with each source image having a corresponding target image with the same name.

#### Stardist training dataset.

DCIS.COM lifeact-RFP cells were incubated for 2h with 0.5 µM SiR-DNA (SiR-Hoechst, Tetu-bio, Cat Number: SC007) before being imaged live for 14h using a spinning-disk confocal microscope (1 picture every 10 min). The spinning-disk confocal microscope used was a Marianas spinning disk imaging system with a Yokogawa CSU-W1 scanning unit on an inverted Zeiss Axio Observer Z1 microscope (Intelligent Imaging Innovations, Inc.) equipped with a 20x (NA 0.8) air, Plan Apochromat objective (Zeiss).

To generate the Stardist training dataset, mask images were generated manually in Fiji. Briefly, the outlines of each nucleus were drawn using the freehands selection tool and added to the ROI manager. Once all outlines were stored in the ROI manager, the LOCI plugin (https://imagej.net/LOCI) was used to create an ROI map. These ROI map images were then used as the mask images to train Stardist.

To automatically track cells, we sequentially used Stardist and the Fiji plugin TrackMate[8]. Briefly, using our Stardist notebook, nuclei from live-cell imaging data were detected, and tracking files containing the coordinate of their center (marked by a dot) were generated (Stardist notebook section 6). These tracking files were then used as input for TrackMate. We also provide in the ZeroCostDL4Mic GitHub page a Fiji macro to batch analyse a folder containing multiple tracking files. Cell tracks were then analysed using the online tool Motility lab (http://www.motilitylab.net/)[9].

**Noise2Void training datasets.**

The 2D dataset provided with our notebooks was generated by plating U-251 glioma cells expressing endogenously tagged paxillin- GFP on fibronectin-coated polyacrylamide gels (stiffness 9.6 kPa)[1]. Cells were then recorded live using a spinning disk confocal microscope equipped with a long working distance of 63x (NA 1.15 water, LD C-Apochromat) objective (Zeiss). The 3D dataset provided with our notebooks was generated by recording A2780 ovarian carcinoma cell, transiently expressing lifeact-RFP (to visualize the actin cytoskeleton), migration on fibroblast-generated cell-derived matrices, further see[10] for methods. The cell-derived matrices were labeled using Alexa Fluor 488 recombinant fibronectin and the images acquired using a spinning disk confocal microscope equipped with a 63x oil (NA 1.4 oil, Plan-Apochromat, M27 with DIC III Prism) objective (Zeiss). For both datasets, the spinning disk confocal microscope used was a Marianas spinning disk imaging system with a Yokogawa CSU-W1 scanning unit on an inverted Zeiss Axio Observer Z1 microscope controlled by SlideBook 6 (Intelligent Imaging Innovations, Inc.). Images were acquired using a Photometrics Evolve, a back-illuminated EMCCD camera (512 x 512 pixels).

**CARE training datasets.**

Briefly, DCIS.COM lifeact-RFP cells were plated on high tolerance glass-bottom dishes (MatTek Corporation, coverslip 1.5) and were allowed to reach confluence. Cells were then fixed and permeabilized simultaneously using a solution of 4% (wt/vol) paraformaldehyde and 0.25% (vol/vol) Triton X-100 for 10 min. Cells were then washed with PBS, quenched using a solution of 1 M glycine for 30 min, and incubated with phalloidin-488 (1/200 in PBS; Cat number: A12379; Thermo Fisher Scientific) at 4°C until imaging (overnight). Just before imaging using SIM, samples were washed three times in PBS and mounted in vectashield (Vectorlabs). The SIM system used was DeltaVision OMX v4 (GE Healthcare Life Sciences) fit-ted with a 60x Plan-Apochromat objective lens, 1.42 NA (immersion oil RI of 1.516) used in SIM illumination mode (five phases and three rotations). Emitted light was collected on a front-illuminated pco.edge sCMOS (pixel size 6.5 µm, read-out speed 95 MHz; PCO AG) controlled by SoftWorx. In the provided dataset, the high signal-to-noise ratio images were acquired from the phalloidin-488 staining using acquisition parameters optimal to obtain high-quality SIM images (in this case, 50 ms exposure time, 10% laser power). In contrast, the low signal-to-noise ratio images were acquired from the LifeAct-RFP channel using acquisition parameters more suitable for live-cell imaging (in this case, 100 ms exposure time, 1% laser power). The dataset provided with the 2D CARE notebooks are maximum intensity projections of the collected data.

**Label-free prediction (fnet) training dataset.**

The dataset provided for training Label-free prediction notebook was designed to predict a mitochondrial marker from brightfield images. Before the acquisition, fixed HeLa ATCC cells were permeabilized and blocked using 0.25% Triton X-100 (Sigma Aldrich, Germany) and 3% IgG-free bovine serum albumin (BSA, Carl Roth, Germany) in PBS for 1.5h. Cells were labeled for TOM20 using 5 µg/ml rabbit anti-TOM20 primary antibody (sc-11415, Santa Cruz, USA) and 10 µg/ml donkey-anti-rabbit-secondary antibody (Alexa Fluor 594 conjugated, A32754, Thermo Fisher, USA) in PBS containing 0.1% Triton X-100 and 1% BSA for 1.5h each. Samples were rinsed twice and washed thrice with PBS (5 min) after each incubation step. Image stacks were acquired on a Leica SP8 confocal microscope (Leica Microsystems, Germany) bearing a 63x, 1.40 NA oil objective (Leica HC PL APO). The pixel size was set to 90 nm in XY-dimensions, and 150 nm in z (32 slices) and fluorescence image stacks were recorded using 561 nm laser excitation and collected by a (photomultiplier tube) PMT. The corresponding transmitted light image stack was recorded in parallel using a transmitted light PMT. We acquired 25 3D stacks with dimensions of 1024x1024x32. To create the training set each image stack was split into four stacks with dimensions of 512x512x32, giving a dataset of 100 images of which 92 were used for training and 8 unseen for testing and quality control (see Table). The raw data were converted into .tif file format and split into stacks of the respective channels (fluorescence and transmitted light). To prepare a training set, stacks were split into individual folders by channel. To create matched training pairs, the signal files (transmitted light) and their respective targets (fluorescence) must be in the same order in the irrespective folders. It is therefore advisable to number source-target pairs or to give the files the same names.

**Deep-STORM training and example dataset.**

For Deep-STORM, training data and test data can be generated via SMLM data simulations that mimic the experimental data type that needs to be subsequently analysed. This is directly possible within the notebook that we provide. The example experimental data that we provide was obtained from a glial cell in a culture of rat hippocampal neurons, fixed and stained with phalloidin-Alexa Fluor 647[11], then imaged on a Nikon N-STORM microscope using 100X, NA 1.49 TIRF objective (256x256 pixels with 160 nm pixel size, acquiring 59,900 frames at 15ms/frame). In order to simulate a higher density of emitters, we binned the data in groups of 4 frames using Fiji (Grouped z-project / Sum slices plugin) leading to 14,975 frames (BIN4), or groups of 10 frames leading to 5,990 frames (BIN10). The binning was performed on a 32-bits dynamic range.

In order to train the model for the reconstruction of the BIN4 and the BIN10 dataset, 20 frames of a 64x64 pixels field-of-view were simulated with parameters matching the experimental datasets closely. The parameter lists are shown in Table 1.

**Table 1**: Simulation parameters used to generate training data for Deep-STORM datasets.

| Dataset | BIN4 | BIN10 | TUB |
|---|---|---|---|
| FOV size (nm) | 10240 | 10240 | 15800 |
| Pixel size (nm) | 160 | 160 | 158 |
| ADC/photon conversion factor | 16 | 16 | 12.7 |
| Readout noise (ADC) | 1040 | 1800 | 370 |
| Offset (ADC) | 10550 | 26500 | 4090 |
| Emitter density (#/um$^2$) | 2.8 | 4 | 3 |
| STD of Emitter density (#/um$^2$) | 0 | 0.8 | 0.5 |
| Number of frames | 20 | 20 | 20 |
| PSF sigma (nm) | 153 | 150 | 160 |
| STD of PSF sigma (nm) | 29 | 25 | 30 |
| Number of | 3500 | 5500 | 2800 |

| | | | |
|---|---|---|---|
| **photons** | | | |
| **STD of Number of photons** | 850 | 2000 | 900 |

The localization files obtained from our Deep-STORM notebook were subsequently imported in ThunderSTORM[12] for drift correction using cross-correlation analysis[13].

For DNA-PAINT imaging of the tubulin cytoskeleton, U2OS cells were fixed and immuno-labeled with mouse-anti-β-tubulin primary antibody (32-2600, ThermoFisher) and DNA-conjugated (P1-docking strand) secondary antibody. DNA-PAINT imaging was performed using a Nikon N-STORM system bearing a Nikon Apo TIRF 100x oil immersion objective (1.49 NA). 2 nM P1-ATTO655 (8 nt duplex, Eurofins) in PBS pH 8.2 + 500 mM NaCl was added to the sample, resulting in a labeling density significantly higher than used for conventional DNA-PAINT imaging. Fluorophores were excited in highly inclined and laminated optical sheet mode using ~1 kW/cm² 647 nm laser illumination. 2,000 frames were recorded at 10 Hz frame rate and an EM gain of 200.

Parameters for simulating the training data (TUB) were estimated from the raw movie or ThunderSTORM ME-MLE reconstruction of a few frames and are listed in Table 1. Training was performed for 30 epochs and 1,750 steps per epoch using a batch size of 4. Both localizations obtained from Deep-STORM and ThunderSTORM analysis were corrected for drift using the cross-correlation function in ThunderSTORM. Emission events that are split onto subsequent frames were merged, applying a 40 nm distance threshold (no dark frame allowed).

### YOLOv2 training dataset.

The YOLOv2 dataset is composed of live-cell movies of breast cancer cells (MDA-MB-231) migrating on cell-derived matrices generated by human fibroblasts[10]. Cell-derived matrices were generated as previously described[14]. MDA-MB-231 cells were seeded at a density of 5,000 cells per ml on cell-derived matrices and allowed to spread for four hours. Cells were then filmed using an inverted widefield microscope (AxioCam MRm camera, EL Plan-Neofluar 20 /0.5 NA objective (Carl Zeiss)) equipped with a heated chamber (37°C) and $CO_2$ controller (5%). Images were collected every 10 min.

To create the annotations, 30 individual images of size 1380x1040 were saved as .png files and loaded into https://www.makesense.ai/, an online annotation platform[15]. A new project was started with the ‘Object Detection’ option. Next, a list of labels was created using the ‘+’ option and typing the names of the classes to be identified. The option ‘Going on my own’ was selected after the creation of the labels and without selecting the ‘COCO SSD...’ or POSE-NET...’ boxes. In the next step, bounding boxes were drawn using the cursor and object classes were added by clicking on the ‘Select Label’ option on the right side of the window and selecting from the previously assembled labels list, now available as a dropdown list. After all images in the dataset were annotated the annotations were downloaded by selecting ‘Export Labels’ and checking the box ‘A .zip package containing the files in VOC.XML format’. The annotation files were then downloaded in a zip folder with the original images’ filenames and the .xml file suffix. These files were used as targets for the training of YOLO network in our notebook. We used the img_aug library[16] to augment the dataset and the bounding boxes by rotation and flipping.

### CycleGAN training dataset.

U2OS cells were plated on fibronectin-coated glass-bottom dishes (MatTek Corporation) for 2h before methanol fixation at -20°C for 5 min. Fixed samples were washed three times using PBS and stained with an anti-tubulin antibody (clone 12G10, Developmental Studies Hybridoma Bank) for 35 min at room temperature. Samples were then washed thrice with PBS and incubated with an anti-mouse secondary antibody conjugated to Alexa488 (ThermoFisher Scientific. Cat num: R37114). Stained samples were washed thrice with PBS and kept at 4°C, in PBS, until imaging.

The SDC dataset was acquired using a Marianas spinning disk equipped with a Yokogawa CSU-W1 scanning unit on an inverted Zeiss

Axio Observer Z1 microscope and a 100× (NA 1.4 oil, Plan-Apochromat, M27) objective and controlled by SlideBook 6 (Intelligent Imaging Innovations, Inc.). Images were acquired using a Photometrics Evolve, back-illuminated EMCCD camera (512 × 512 pixels). For each field of view, 200 images were acquired. The SDC images used to train CycleGAN were generated by performing average projections of the collected images. To maintain a uniform pixel size across the SDC, SIM and FBSR images, the SDC images were magnified by 4 using a bilinear interpolation.

The fluctuation-based super-resolution dataset was acquired by processing the SDC images using the latest implementation of NanoJ-SRRF[17] within the ImageJ software[7]. This new version of SRRF is available upon request and will be openly available for download soon. The SRRF settings were chosen so that the least amount of errors were present in the reconstructed images (estimated using SQUIRREL[18]) and were as followed: "vibration correction", on; Radius, 2; Sensitivity, 2; magnification, 4; temporal analysis, average; intensity weighting, on; macro-pixel patterning correction, on.

The SIM dataset was acquired using a DeltaVision OMX v4 (GE Healthcare Life Sciences) fitted with a 60x Plan-Apochromat objective lens, 1.42 NA (immersion oil RI of 1.512) used in SIM illumination mode (five phases and three rotations). Emitted light was collected on a front-illuminated pco.edge sCMOS (pixel size 6.5 um, readout speed 95 MHz; PCO AG) controlled by SoftWorx. Each dataset was augmented by five by randomly cropping (original size 2048x2048 px, crop size 1280x1280), flipping and rotating the original images. This augmentation pipeline was generated using Augmentor[19], for which we also provide a ZeroCostDL4Mic notebook.

**pix2pix training dataset.**

DCIS.COM lifeact-RFP cells were incubated for 2h with 0.5 µM SiR-DNA (SiR-Hoechst, Tetu-bio, Cat Number: SC007) before being imaged live for 14h (1 picture every 10 min) using a spinning-disk confocal microscope. The spinning-disk confocal microscope used was a Marianas spinning disk imaging system with a Yokogawa CSU-W1 scanning unit on an inverted Zeiss Axio Observer Z1 microscope (Intelligent Imaging Innovations, Inc.) equipped with a 20x (NA 0.8) air, Plan Apochromat objective (Zeiss).
