## Supplementary Information for "ZeroCostDL4Mic: an open platform to use Deep-Learning in Microscopy"

Lucas von Chamier[1] *, Romain F. Laine[1,2] *, Johanna Jukkala[3,4], Christoph Spahn[5], Daniel Krentzel[6,7], Elias Nehme[8,9], Martina Lerche[3], Sara Hernández-Pérez[3,10], Pieta K. Mattila[3,10], Eleni Karinou[11], Séamus Holden[11], Ahmet Can Solak[12], Alexander Krull[13-15], Tim-Oliver Buchholz[13,14], Martin L. Jones[6], Loïc A Royer[12], Christophe Leterrier[16], Yoav Shechtman[9], Florian Jug[13,14,17], Mike Heilemann[5], Guillaume Jacquemet[3,4], and Ricardo Henriques[1,2,18]

[1]MRC-Laboratory for Molecular Cell Biology, University College London, London, UK

[2]The Francis Crick Institute, London, UK

[3]Turku Bioscience Centre, University of Turku and Åbo Akademi University, Turku, Finland

[4]Faculty of Science and Engineering, Cell Biology, Åbo Akademi University, Turku, Finland

[5]Institute of Physical and Theoretical Chemistry, Goethe-University Frankfurt, Frankfurt, Germany

[6]Electron Microscopy Science Technology Platform, The Francis Crick Institute, 1 Midland Road, London NW1 1AT, UK

[7]Department of Bioengineering, Imperial College London, South Kensington Campus, London, SW7 2AZ, UK

[8]Department of Electrical Engineering, Technion - Israel Institute of Technology, Haifa, Israel

[9]Department of Biomedical Engineering, Technion - Israel Institute of Technology, Haifa, Israel

[10]Institute of Biomedicine, and MediCity Research Laboratories, University of Turku, Finland

[11]Centre for Bacterial Cell Biology, Biosciences Institute, Faculty of Medical Sciences, Newcastle University, UK

[12]Chan Zuckerberg Biohub, San Francisco, CA, USA

[13]Center for Systems Biology Dresden (CSBD), Dresden, Germany

[14]Max Planck Institute for Molecular Cell Biology and Genetics, Dresden, Germany

[15]Max Planck Institute for Physics of Complex Systems, Dresden, Germany

[16]Aix Marseille Université, CNRS, INP UMR7051, NeuroCyto, Marseille, France

[17]Fondatione Human Technopole, Milano, Italy

[18]Instituto Gulbenkian de Ciência, Oeiras, Portugal

*Equal contributing authors

**Supplementary Information: Contents**

**Supplementary Note 1: When in doubt, always retrain! A supplementary discussion**

**Supplementary Note 2: Image analysis tasks currently implemented in ZeroCostDL4Mic**

**Supplementary Note 3: Quality Control on trained models**

**Supplementary Note 4: Data augmentation**

**Supplementary Note 5: Transfer Learning and training from a previously saved model**

**Supplementary Note 6: Capabilities of Google Colab**

**Supplementary Note 7: Future perspectives for ZeroCostDL4Mic**

**Supplementary Table 1. Overview of the available datasets used for training the networks.**

**Supplementary Table 2. Hyperparameters used to train the networks, GPU types allocated, and corresponding training times.**

**Supplementary Fig. 1. Of the importance of training models on suitable data.**

**Supplementary Fig. 2: Image segmentation networks (U-Net and StarDist).**

**Supplementary Fig. 3: Object detection (YOLOv2).**

**Supplementary Fig. 4: Image denoising and restoration networks (CARE and Noise2Void).**

**Supplementary Fig. 5: Super-resolution microscopy network (Deep-STORM).**

**Supplementary Fig. 6: Image-to-image translation networks (fnet, pix2pix and CycleGAN).**

**Supplementary Fig. 7: Example illustrating how ZeroCostDL4Mic notebook can be used together.**

**Supplementary Fig. 8: Graphical user interface (GUI) of the ZeroCostDL4Mic notebooks.**

**Supplementary Fig. 9: Quality Control of trained models.**

**Supplementary Fig. 10: Quality Control in the ZeroCostDL4Mic notebooks.**

**Supplementary Fig. 11. Model performance vs training parameters.**

**Supplementary Fig. 12: Example of data augmentation section in the ZeroCostDL4Mic fnet notebook.**

**Supplementary Fig. 13: Data augmentation improves prediction performance.**

**Supplementary Fig. 14: Transfer learning in the ZeroCostDL4Mic notebooks.**

**Supplementary Fig. 15: Transfer learning can reduce training time and improve performance**.

**Supplementary Video 1**: Full run-through of the workflow to obtain the notebooks and the provided test datasets as well as a common use of the notebook. YouTube link: https://youtu.be/TrDuidvO85s

**Supplementary Video 2**: Representative results obtained from the provided test dataset. YouTube link: https://youtu.be/KauKEr0Kkkc

**Supplementary Video 3:** Video highlighting how a 3D U-Net network trained using ZeroCostDL4Mic can be used to segment electron microscopy data. The raw data, as well as the training target (magenta) and the model predictions (green), are displayed. The visualization of the 3D reconstruction was generated using Imaris. YouTube link: https://youtu.be/gkFOv4k3eQU

**Supplementary Video 4**: Video highlighting how a StarDist model, trained using ZeroCostDL4Mic, can be used in conjunction with TrackMate[1] to track cell movement automatically. Raw data (Input), StarDist predictions as well as local cell tracks are displayed. Cells are DCIS.COM cells labeled with SiR-DNA and imaged using a spinning disk confocal microscope. YouTube link: https://youtu.be/poYoDer9Pj0

**Supplementary Video 5**: Video highlighting how a YOLOv2 model, trained using ZeroCostDL4Mic, can be used to detect and classify cell shapes in brightfield time-lapse data of MDA-MB-231 cells migrating on cell-derived matrices. YouTube link: https://youtu.be/J6Ec8WVojYk

**Supplementary Video 6:** Example highlighting how a CARE network, trained using ZeroCostDL4Mic, can be used to denoise SIM live-cell imaging data. Here, a 3D CARE network was trained using SIM images of the actin cytoskeleton of DCIS.COM cells using fixed samples to denoise live-cell imaging data (see Online methods for details). Both input and the model predictions are displayed (representative single z plane). YouTube link: https://youtu.be/IWt9eQMmseE

**Supplementary Video 7:** Example highlighting how a Noise2Void 3D network, trained using ZeroCostDL4Mic, can be used to denoise live-cell imaging data. Movie of an ovarian carcinoma cell labeled with lifeact-RFP migrating on cell-derived matrices (labeled for fibronectin). Images were acquired using a spinning disk confocal microscope. YouTube link: https://youtu.be/joEsIB4aH18

**Supplementary Video 8**: Example highlighting how a Noise2Void 3D network, trained using ZeroCostDL4Mic, can be used to denoise microscopy data. Dataset #1: Live HeLa cells labeled with MitoTracker Red CMX Ros imaged with a VisiTech iSIM microscope, each frame represents a 20s interval. Dataset #2: Fixed HeLa cells immuno-labeled for microtubules and mitochondria imaged using a Leica SP8 confocal microscope. YouTube link: https://youtu.be/D_IC_nHJGI8

**Supplementary Video 9**: Example highlighting how a Noise2Void 2D network, trained using ZeroCostDL4Mic, can be used to denoise live-cell imaging data. Video of a glioma cell endogenously labeled for paxillin-GFP, migrating on 9.6 kPa polyacrylamide hydrogel. Images were acquired using a spinning disk confocal microscope. Both the training source and model predictions are displayed. YouTube link: https://youtu.be/khHHQQynsfg

**Supplementary Video 10**: Example highlighting how a Deep-STORM network, trained using ZeroCostDL4Mic, can be used to reconstruct high-density SMLM data. Reconstruction of a glial cell labeled with Phalloidin-AlexaFluor 647 (BIN4 dataset). The video shows the raw data, the wide-field equivalent data (obtained from binning the drift-corrected raw data), and the Deep-STORM reconstruction. YouTube link: https://youtu.be/fk3P9Kei3Us

**Supplementary Video 11:** Example showcasing how ZeroCostDL4Mic notebooks can be used sequentially. Here we wanted to automatically track the migration pattern of DCIS.COM cells labeled with lifeact-RFP. Therefore, we first used pix2pix to “translate” the actin staining into nuclei staining and StarDist to detect the nuclei. From the StarDist prediction, cells were tracked automatically using TrackMate. In the video, the raw data (lifeact-RFP), the pix2pix predictions (Fake SiR-DNA), the StarDist predictions, and the TrackMate tracks are displayed. YouTube link: https://youtu.be/VtP9QjJjxP0

**Supplementary Note 1: When in doubt, always retrain! A supplementary discussion**

The primary focus of ZeroCostDL4Mic is to provide a straightforward and free platform to aid novice users in using Deep Learning (DL) in microscopy. A vital component of this platform is the capacity to simplify model training, which remains a significant difficulty. Because it can be challenging to train DL networks (in time, resources, and skills), several labs are taking the approach of providing pre-trained network models, which can then be used to process imaging data[2–5].

Pre-trained models can be useful as they can be easily re-used to analyze data that is very similar to the one used during the initial training. However, pre-trained models should be used with caution on new data as they tend to be very specific to the microscopes and samples used to generate the training dataset. The inappropriate use of pre-trained models can lead to erroneous results when applied to a different dataset type[6,7], which, unfortunately, may lead to visually pleasing yet inaccurate results[8,9]. For example, Supplementary Fig. 1 shows how using an inappropriate model can lead to erroneous results in the prediction. Specifically, a CARE 3D network was trained to denoise either actin (lifeact) or mitochondria (TOM20) data and used to denoise both types of datasets. When using the incorrect model (a model trained on actin to restore mitochondria and vice-versa), the predictions present artifacts and significantly weaker quality control metrics (see Supplementary Note 3 for details on quality control metrics).

Given this issue, it becomes critical for researchers to have the option to train models (or re-train models, using transfer learning, see Supplementary Note 5 dedicated to transfer learning) using their specific data of interest to produce high-fidelity and reliable results.

ZeroCostDL4Mic allows researchers to access the resources needed to train or re-train using transfer learning a range of DL networks in an accessible and educational manner. It provides an easy-to-follow user interface (see Supplementary Fig. 8), allowing for the notebook code to be hidden by default. It is notwithstanding easy to show and edit the underlying network code via the notebook page itself (by double-clicking on the cell of interest). This option allows researchers to explore the code and to interactively test and learn the programmatic basis of each code section, if necessary. ZeroCostDL4Mic offers a unique and powerful way for users to explore the code structure and concepts and build more confidence and understanding in the analysis workflow. We believe this platform takes a significant step to increase the uptake and the understanding of DL tools by microscopy users and highlights the need for all DL users to grasp and tackle the limitations of the approach.

**Supplementary Note 2: Image analysis tasks currently implemented in ZeroCostDL4Mic**

Within the ZeroCostDL4Mic platform, we implemented several important bioimage analysis tasks. We grouped the implemented networks in the four following general tasks:

- Image segmentation and object detection
- Image restoration (denoising)
- Super-resolution microscopy
- Image-to-image translation

### 2.1 Image segmentation and object detection

Image segmentation and object detection are related tasks that both identify specific image areas of interest and highlight them against the background. However, image segmentation and object detection can be distinguished into subtly different tasks, segmentation being more concerned with the identification of objects' boundaries and object detection with the presence and identity of various objects in the image. Typically, segmentation extracts image masks while object detection identifies objects using bounding boxes, delimiting the extent of the detected object within the image.

#### 2.1.1 Image segmentation

Manual segmentation is a time-intensive and challenging task that typically requires expert knowledge. Segmentation is often a significant bottleneck for studies that aim at quantifying large datasets[10,11]. Hence, it has become a field where DL is of great interest since it can combine expert-level performance with high-throughput analysis[12]. Within ZeroCostDL4Mic, we implemented two networks, U-Net and StarDist, to perform segmentation tasks.

U-Net is a generalist DL network based on an encoder-decoder architecture[13,14]. U-Net was initially developed for the segmentation of EM images, although it has since been adapted for many other tasks. In particular, a number of the other DL networks provided within ZeroCostDL4Mic are derived from or inspired by U-Net (e.g., CARE, StarDist 2D, Noise2Void, fnet and Deep-STORM). To illustrate the applicability of our ZeroCostDL4Mic U-Net notebooks, we trained a U-Net network to segment membranes from 2D electron microscopy (EM) images (Supplementary Fig. 2a). The training dataset used here is publicly available (ISBI 2012 segmentation challenge dataset[15]) and is composed of 2D EM images and associated masks.

Once trained using ZeroCostDL4Mic, a U-Net model provides segmentation probability maps that need to be thresholded to obtain binary masks (Supplementary Fig. 2a). In our notebooks, the optimal threshold can be estimated using the quality control section (see Supplementary Note 3 for details on quality control metrics) (Supplementary Fig. 2a). To further demonstrate the versatility of the U-Net architecture, we next trained, using ZeroCostDL4Mic, a 3D U-Net model to segment mitochondria from 3D EM images (5x5x5 $\mu m^3$ section taken from the CA1 hippocampus region of the brain, Supplementary Fig. 2b). For this, we used another publicly available segmentation dataset (from EPFL: https://www.epfl.ch/labs/cvlab/data/data-em/)[16]. The 3D mitochondria segmentation obtained using our notebook is in excellent agreement with the ground truth data for this test dataset (Supplementary Fig. 2b and Supplementary Video 3).

StarDist is a DL method optimized to segment nuclei in 2D or 3D microscopy images[17,18]. Training a StarDist network requires the user to provide matching images of nuclei and their corresponding masks. To showcase our ZeroCostDL4Mic notebooks, we first trained a StarDist model to segment the nuclei of densely packed cell monolayers (Supplementary Fig. 2c). The StarDist model, trained with ZeroCostDL4Mic, enabled almost perfect segmentation of these images. Interestingly, this model was able to detect extra nuclei that were missed when these images were manually labeled (Supplementary Fig. 2c). To further showcase the functionality of our StarDist notebooks, we next used a StarDist model, trained using ZeroCostDL4Mic, to analyze live-cell imaging data (Supplementary Fig. 2d). To simplify the automated tracking of StarDist detected nuclei, our StarDist 2D notebook generates a file containing the coordinates of all the nuclei in the images, called tracking files, that can be used as input for TrackMate, a popular Fiji[19] tracking plugin[11]. By combining automated nuclei segmentation (using StarDist) and automatic tracking (using TrackMate), our ZeroCostDL4Mic StarDist notebook enables automated cell tracking (Supplementary Fig. 2d and Supplementary Video 4). In our ZeroCostDL4Mic GitHub repository, we also provide a Fiji macro to batch analyze a folder containing multiple tracking files.

#### 2.1.2 Object detection

Object detection is a popular discipline in DL challenges[20,21] since it is highly applicable to real-life cases such as self-driving cars[22]. It has also become of interest in microscopy studies that need to identify classes or subsets of objects in an image, e.g., counting pathogens or identifying cell types in images[23,24].

To provide object detection capabilities within the ZeroCostDL4Mic platform, we implemented the DL network YOLOv2[25]. YOLOv2 learns to detect objects within an image by identifying bounding boxes encasing the detected objects and classifying the object within each box. This detection and classification are performed by supervised learning from annotated training datasets.

YOLOv2 continues to be considered one of the best networks for such tasks and is commonly used in DL challenges[26,27]. To demonstrate how our YOLOv2 implementation can be applied to microscopy data, we acquired time-lapses of MDA-MB-231 cells migrating on cell-derived matrices using brightfield microscopy. The YOLOv2 network in our ZeroCostDL4Mic notebook was trained with input images, which were respectively paired with a file containing the bounding box coordinates and class labels of the objects in the image (generated using the web tool makesense.ai[28]). In this example, we annotated our dataset using four different classes, according to cell shape: "*elongated*" cells, "*rounded*" cells, "*dividing*" cells and "*spread-out*" cells (see Supplementary Fig. 3a). Once trained the model can be applied to unseen data and our YOLOv2 notebook provides images with bounding boxes (Supplementary Fig. 3b) and a .csv file which contains the bounding box coordinates and classes of all identified objects. This .csv file can be imported to Fiji as a results table and used in the ROI manager for further analysis. The network, trained using ZeroCostDL4Mic, correctly classified and labeled many cells in the image (as indicated by the mAP score, see Supplementary Note 3 for details about quality control metrics). Our notebook also accommodates batch processing, which is especially useful for the analysis of time-lapse datasets. (Supplementary Video 5).

### 2.2 Image restoration (denoising)

Fluorescence live-cell imaging has become the primary strategy to observe molecular-specific biological structures and pathways in living organisms. However, when performing live-cell imaging experiments, typically, low laser intensities need to be used to ensure the survival of the biological specimen observed[29]. Also, to improve the observations' physiological relevance, it is advisable to keep the expression level of the molecule of interest as close as possible to the endogenous levels. As most proteins are expressed at low levels in cells, the amount of fluorescence signal collected during live microscopy experiments is often meager. Because of these two factors, live-cell imaging experiments often lead to the acquisition of noisy images, and denoising is becoming increasingly essential for interpretation of the data and further analysis. Within ZeroCostDL4Mic, we implemented two networks, CARE[3] and Noise2Void[30], specialized in the denoising of microscopy data.

Content-aware image restoration (CARE)[3] is a DL method capable of image restoration from corrupted bio-images (whether corruption comes from noise, artifacts, or low resolution, for instance).

The network allows image denoising and resolution improvement in 2D and 3D images, using supervised training.

To train a CARE network, the user needs to provide a training dataset made of paired images, such as low signal-to-noise ratio (SNR) images and high SNR images of the same fields-of-view.
However, the specific function of the network is determined by the images provided in the training dataset. For instance, if noisy images are provided as input, and high SNR images are provided as targets, the network will perform denoising. To illustrate the applicability of our ZeroCostDL4Mic CARE notebooks, we trained a 3D CARE network to denoise live-cell structured illumination microscopy (SIM) imaging data (Supplementary Fig. 4a and 4b). To generate a suitable training dataset, both high and low SNR images of the actin cytoskeleton were acquired from fixed samples. Specifically, high SNR images were acquired by staining lifeact-RFP DCIS.COM cells with phalloidin-488 and using acquisition parameters optimal to obtain high-quality SIM reconstructions. In contrast, the low SNR images were acquired from the lifeact-RFP channel using acquisition parameters typically used for live-cell SIM imaging (see Online Methods for acquisition details). The network trained using these images was then used to restore live-cell imaging data (Supplementary Fig. 4b, Supplementary Video 6). The approach employed here is especially useful as it can be very challenging to obtain high-quality live-cell imaging, otherwise using SIM for extended periods of time[7,31]. We also noted that this approach somewhat helped suppress some typical patterns originating from low SNR SIM reconstructions.

Noise2Void[30] is a DL method that was designed to perform denoising of microscopy images in the absence of a dedicated paired training dataset. In other words, Noise2Void can learn how to denoise a dataset directly from the dataset to be denoised. Therefore, no specific training datasets are required, only noisy images. This is the main advantage of Noise2Void compared to CARE. However, it also means that it can be more challenging to assess the trained model's quality as ground-truth images may not be readily available. Here, we provide several examples of data that can be generated using the ZeroCostDL4Mic Noise2Void notebooks (Supplementary Fig. 4c and 4d). In the first example, we used our Noise2Void notebook to denoise the movie of an ovarian carcinoma cell, labeled with lifeact-RFP, migrating on cell-derived matrices (Supplementary Fig. 4c). In this case, a Noise2Void model was trained with each fluorescence channel independently. In both cases, a single Z stack (time-point) was used to train Noise2Void, and the resulting model was applied to the rest of the movie (Supplementary Fig. 4c and Supplementary Video 7).

We also demonstrate the applicability of our Noise2Void notebook to train networks to denoise time-lapse and multi-color fluorescence microscopy data acquired using different types of microscope. The time-lapse dataset was obtained from live cells labeled with MitoTracker Red CMX Ros and imaged using an iSIM microscope.

The dual-color fluorescence dataset was obtained from fixed samples immuno-labeled to visualize microtubules and mitochondria and imaged using a confocal microscope (Supplementary Video 8). Also, in Supplementary Fig. 4d, we showcase how our Noise2Void notebook can be used to improve low-signal fluorescence imaging, by denoising data capturing the endogenous expression levels of a glioma cell endogenously labeled for paxillin-GFP, migrating on polyacrylamide hydrogel (see also Supplementary Video 9).

In all the displayed examples, the Noise2Void models trained in ZeroCostDL4Mic performed very well and significantly improved the images' quality. However, it is important to note that, in our hands, Noise2Void did not perform well when used to denoise the actin dataset used to train CARE (Supplementary Fig. 4a). This is likely because the noise in that particular dataset is not homogeneous and contains structures (from the SIM reconstruction process). This highlights the importance of testing a range of networks to identify which approach is most suited for a specific dataset, underlying the utility of a platform integrating a large range of networks.

#### 2.3 Super-resolution microscopy

Within ZeroCostDL4Mic, we also implemented DL-enabled super-resolution microscopy by integrating the Deep-STORM[32] network. Deep-STORM is a neural network capable of reconstructing single-molecule localization microscopy (SMLM) data to obtain super-resolution images from relatively dense emitter datasets. Performing high-density SMLM reconstruction has the advantage of achieving high-performance SMLM imaging in poorly blinking conditions and offers the possibility of significantly shortening the image acquisitions. Here, the network can learn from raw SMLM data accompanied by ground truth localization coordinates to output a sparse image where each localization is displayed as a single pixel in an upsampled image. The ZeroCostDL4Mic implementation of Deep-STORM also offers the additional feature of extracting localization coordinates from this upsampled image, enabling further analysis such as drift correction (also directly possible within the notebook) or spatial point pattern analysis. One key advantage of Deep-STORM is that it can be fully trained on simulated data, so we included a simulator within the notebook to directly generate SMLM training (and test) data that can be tuned to mimic the experimental data type that needs to be subsequently analyzed.

Supplementary Fig. 5 shows representative reconstructions obtained from Deep-STORM from relatively dense emitter SMLM data (as shown in the first column, first and second row of Supplementary Fig. 5 and in Supplementary Video 10).

We show two different datasets: actin labeling via phalloidin-AlexaFluor 647 *d*STORM imaging (Supplementary Fig. 5a, BIN10 data, see Online methods for details) and DNA-PAINT[33] tubulin immuno-labeling via a P1-ATTO655 imager strand (Supplementary Fig. 5b). For comparison, we processed the same data using Multi-Emitter Maximum Likelihood Estimation (ME-MLE) using ThunderSTORM[34]. Deep-STORM led to high-quality reconstruction in a fraction of the time necessary for ME-MLE to produce the reconstructed image. Additionally, a SQUIRREL[35] analysis of these reconstructions (Supplementary Fig. 5c and 5d) with respect to the equivalent wide-field image (obtained by binning the drift-corrected raw data image stack) allowed us to show that the results obtained from Deep-STORM showed a better agreement with the equivalent wide-field image, highlighting a better linearity of the reconstructions compared to that obtained with ME-MLE.

#### 2.4 Image-to-image translation

The image-to-image translation task refers to DL networks that can transform one type of image into another (for instance, predict a fluorescent label from brightfield images or predict a fluorescent label from another fluorescent label). Within ZeroCostDL4Mic, we implemented three networks, label-free prediction (fnet)[36], pix2pix[37], and CycleGAN[38] capable of performing image-to-image translations. Here, we introduce these networks depending on the type of training data required to train them.

##### 2.4.1 Image-to-image translation requiring a paired training dataset

Image-to-image translation networks such as fnet and pix2pix require the user to provide a paired dataset for training as they rely on a fully supervised training approach to learn the transformation. This means that the same field-of-view needs to be acquired in the two experimental conditions and be provided with an indication of correspondence (Supplementary Fig. 6a).

fnet[36] is a DL method developed as a tool for label-free predictions from unannotated brightfield and EM images. fnet utilizes an encoder-decoder architecture based on U-Net and training requires paired 3D stacks of two channels, e.g. fluorescence and transmitted light. To showcase the ZeroCostDL4Mic fnet notebook, we trained a fnet model to predict TOM20 mitochondrial labeling from brightfield images (Supplementary Fig. 6b).

pix2pix[37] is a DL method that uses generative adversarial networks (GAN)[39] to translate one type of image into another. It is often used to transform drawings into somewhat realistic images of animals (https://affinelayer.com/pixsrv/index.html?ref=producthunt). To train, pix2pix requires paired 2D images. While our pix2pix ZeroCostDL4Mic notebook can potentially be used for any kind of image-to-image translation, we demonstrate a possible biological application for pix2pix by training it to convert the fluorescence image of one label (actin) into that of another label (nucleus) in migrating DCIS.COM cells (Supplementary Fig. 6c).

#### 2.4.2 Image-to-image translation requiring only unpaired training dataset

Image-to-image translation networks such as CycleGAN[38] can capture the characteristics of one image domain and understand how these characteristics could be translated into another image domain, all in the absence of any paired training examples (Supplementary Fig. 6d). This approach, therefore, corresponds to an unsupervised training scheme. As for pix2pix, CycleGAN uses GANs to perform the image-to-image translation task. Not requiring a paired training dataset is the main advantage of CycleGAN compared to pix2pix, which releases constraints in generating paired training dataset. However, it also means that it is more challenging to assess the trained model's quality as paired training sources and training targets are not readily available. While CycleGAN can potentially be used for any image-to-image translation[40], we demonstrate here that it can be used to predict what a fluorescent label would look like when imaged using other imaging modalities. In particular, we trained CycleGAN to predict what images of microtubules acquired with a spinning disk confocal would look like when processed with SRRF[41,42] or imaged with a SIM microscope (Supplementary Fig. 6e and 6f). Using our notebook, we also trained CycleGAN to transform SRRF images into SIM images (Supplementary Fig. 6g).

### 2.5 ZeroCostDL4Mic within larger analysis pipeline

While ZeroCostDL4Mic notebooks are self-contained (they allow users to train/re-train, assess and use DL models), they can also be implemented within larger analysis pipelines. For instance, we demonstrate that data generated by ZeroCostDL4Mic notebooks can significantly facilitate further analysis by seamlessly integrating our StarDist notebook with the Fiji plugin TrackMate[1] and analyze cell migration behavior (Supplementary Fig. 2d). As another example, results generated by our Deep-STORM notebook can be processed with the Fiji plugin ThunderSTORM (Supplementary Fig. 5). We took the approach of making sure that the networks' output is directly compatible with commonly-used software independently of the coding language used.

This allowed us to focus on developing ZeroCostDL4Mic as a Python-based package, ideally suited for DL while making use of pre-existing software developed in Java such as ImageJ/Fiji[19].

Another key point is that ZeroCostDL4Mic notebooks can also easily be used sequentially to perform successive tasks and build an analysis pipeline. To illustrate this, we performed the automatic tracking of migration patterns of DCIS.COM cells labeled with lifeact-RFP. This type of image analysis can be challenging as these cells migrate collectively and are hard to segment using only an actin staining. Therefore, we first used pix2pix to predict a nuclear marker (here predicting a SiR-DNA staining) directly from the actin fluorescent label. The individual nuclei predicted this way were then identified and segmented using StarDist, and cells were automatically tracked using TrackMate (Supplementary Fig. 7 and Supplementary Video 11).

**Supplementary Note 3: Quality control of trained models**

The reliable implementation of DL methods depends on a careful evaluation of the models' output performance; we call this step quality control (QC). QC is crucial to avoid using models that produce low-quality images and artifacts, especially when they may not be easily identifiable by simple visual assessment. ZeroCostDL4Mic implements QC on all the trained models (Supplementary Fig. 9). These metrics can thus help the user improve the models created by comparing models trained with different hyperparameters or explore the range of applicability to different data from which it was trained on (generalization). All the notebooks we provide contain a section dedicated to QC, evaluating the performance of trained models (Supplementary Fig. 10). This section typically has two parts:

- Inspection of the loss function over the number of epochs trained.
- Evaluation of image quality metrics by comparing the model predictions against a ground truth equivalent.

The first performance metrics shown to users are loss curves for model training and validation (Supplementary Fig. 9a). They allow users to determine if the model is overfitting during training, identifiable by an increasing divergence between the validation and training loss. This divergence appears if the model learns features specific to the training dataset instead of general features applicable to all similar datasets, therefore preventing it from generalizing to unseen data, a common problem for deep neural networks[8].

However, even a model that fits well to the validation data can produce unwanted results when used on unseen data, ultimately making the model unreliable. These issues are identifiable by comparing the predictions from unseen data to the equivalent ground-truth data. The metrics used in individual notebooks vary to reflect the differences in the type of data these models operate with. For networks producing a grey-scale image, e.g. CARE, Noise2Void, U-Net, and Label-free prediction (fnet), the metrics used are SSIM (Structural similarity)[43] and RSE (Root Square Error) (Supplementary Fig. 9b). For networks producing a binary or semantic image, e.g., StarDist, the metric used is IoU (intersection over union) (Supplementary Fig. 9c). The YOLOv2 notebook uses the mean average precision score (mAP) as in Everingham *et al.*[21], reflecting the validity of the bounding box positions and the corresponding classification. Below, we describe these metrics and our implementations in detail.

The author of CycleGAN and pix2pix does not recommend the visual inspection of the loss function curves to evaluate the training quality achieved with these networks. Therefore, these training curves are not displayed in the pix2pix and CycleGAN notebooks.

Instead, these two notebooks save model checkpoints every five epochs; the QC section helps to identify the best checkpoint to use and retrieves the corresponding optimal model. Specifically, these QC sections allow the user to perform predictions using all the saved checkpoints and estimate the quality of these predictions by comparing them to the provided ground truth images using the SSIM metrics.

#### 3.1 Structural Similarity Index (SSIM)

First introduced by Wang *et al.*[43] SSIM is calculated as follows:

$$SSIM(X,Y) = \frac{(2\mu_x\mu_y + C_1)(2\sigma_{xy} + C_2)}{(\mu_x^2 + \mu_y^2 + C_1)(\sigma_x^2 + \sigma_y^2 + C_2)}$$

where X and Y are the images to be compared, $\mu_x$ and $\mu_y$ the mean pixel values, $\sigma_x^2$ and $\sigma_y^2$ the variance of pixel values and $\sigma_{xy}$ the covariance between the pixel values in the images. $C_1$ and $C_2$ are constants introduced to avoid instability for small denominators and are defined as:

$$C_1 = (K_1 L)^2$$

$$C_2 = (K_2 L)^2$$

where L is the bit-depth of the images to be compared (for 16-bit images used for the quality assessment in the notebooks $L = 65{,}536$) and $K_1 = 0.01$ and $K_2 = 0.03$, as suggested by the authors of SSIM[43].

To calculate the SSIM on a consistent dynamic range, images are normalized to values between 0 and 1, first by percentile normalization on all data (source, target, and prediction). Then, both source and prediction are further normalized by linear regression compared to the target.

The percentile normalization was performed as follows:

$$I_{ij}^{norm} = \frac{I_{ij} - I_{99.9}}{I_{99.9} - I_{0.1}}$$

where $I_{ij}^{norm}$ represents the normalized intensity value at pixel *(i,j)*, $I_{ij}$ the intensity to be rescaled, $I_{99.9}$ the value of the pixel which lies in the 99.9th percentile of pixel values in the image and $I_{0.1}$ the pixel value of the pixel in 0.1th percentile of pixel values in the image. This percentile-based normalization (instead of using minimal and maximal pixel values) is aimed to prevent the influence of *dead* or *hot* pixels which are common in microscopy images and may distort the useful dynamic range of the image upon normalization.

The linear regression normalization is done the same way as in CARE[3], based on least square minimization, defined as:

$$(\alpha_o, \beta_o) = argmin_{\alpha,\beta} \sum_{i,j} \left( GT_{ij} - \left( \alpha I_{ij} + \beta \right) \right)^2$$

where $GT$ is the ground truth image, $I$ the predicted image, $GT_{ij}$ and $I_{ij}$ the respective pixels in the ground truth and prediction and α and β the parameters which rescale the prediction to the dynamic range of the ground-truth.

The normalized image is then calculated as follows:

$$I^{norm} = \alpha_o I + \beta_o$$

After normalization, the pixel values above 1 and below 0 are thresholded to 1 and 0, respectively.

Values for SSIM can vary between -1 and 1, with 1 indicating a perfect structural content agreement between the two images. In practice, SSIM values rarely fall below 0, which would represent inverted structures in the image. For this reason, we truncate SSIM values to a range between 0 and 1. The SSIM map is calculated on a local window around the pixel of interest. The window size is set to 11x11 pixels and a Gaussian weighting function of 1.5 pixels standard deviation. The similarity map obtained displays areas with high and low similarity in different colors, enabling inspection of areas in the image where the model performs better or worse. Therefore, a perfectly-performing model will produce an SSIM map with nearly uniform values close to 1 across the image. A global SSIM metric can also be estimated over the whole image (mSSIM). This is simply determined by calculating the average SSIM value over the whole SSIM map. This value allows an overall quantitative comparison of performance between different images or network parameters.

In the notebook, the SSIM map is calculated between ground-truth against model prediction and between ground-truth against source image (when appropriate). This way, the metric can be used comparatively to judge the improvement of the prediction over the source image.

The use of SSIM for QC is demonstrated in Supplementary Fig. 9b, where we compare the prediction of a CARE model trained to denoise fluorescence images of mitochondria (TOM20), with its expected denoised ground-truth. The almost uniform SSIM map and an mSSIM index above 0.9 signify a high level of similarity between the prediction and the ground-truth. It also shows a clear improvement to the SSIM map and index calculated between ground truth and the noisy input image, suggesting that the network improved the input image.

#### 3.2. Root Squared Error (RSE)

The RSE is obtained by calculating the absolute difference between two images on a pixel-per-pixel basis. By calculating the RSE between the prediction and the target image (ground-truth), this error map shows areas of high and low errors in different colors, providing a second metric for inspecting local artifacts.

$$RSE_{ij} = \sqrt{\left(GT_{ij} - I_{ij}^{norm}\right)^2}$$

Similarly to SSIM, we also provide an image-wise metric as a normalized root mean squared error (*NRMSE*) between the images defined as:

$$NRMSE = \sqrt{\sum_{i,j}\left(GT_{ij} - I_{ij}^{norm}\right)^2}$$

The RSE map and NRMSE reflect the amount of discrepancy between two images. In this case, good performance is indicated by low values for these metrics. A perfect agreement with the ground truth image will lead to RSE = 0 across the image and NMRSE = 0.

In the notebooks using RSE, a comparison between prediction against ground-truth and source against ground-truth is shown side-by-side to indicate how much the image has improved (i.e. brought closer to the ground-truth than the source) by the trained model. The output of the RSE metric is also shown in Supplementary Fig. 9b, where we compare the prediction of the trained CARE network with the expected ground-truth. The RSE map is almost uniformly dark and the NRMSE score is low, suggesting a good match between ground truth and prediction. The comparison with the same metrics calculated between the noisy input and the ground truth also shows how the CARE prediction improved the image by reducing the errors seen in the RSE map and by lowering the value of the NRMSE compared to the input image.

#### 3.3 Intersection over union (IoU)

For the U-Net and StarDist segmentation networks, we use the intersection over union (IoU) metric, commonly used for segmentation performance evaluation[17]. To test how well the model segments an input image the ground-truth mask is compared to the predicted segmentation mask by dividing the number of pixels shared between both masks by the total number of pixels in the union of the two masks:

$$IoU = \frac{I \cap GT}{I \cup GT}$$

where $I$ represents the predicted image and $GT$ the ground-truth image.

The IoU metric quantifies the percentage of overlap between the target mask and the prediction output. Therefore, scores closer to 1 typically mean better performance of the trained model.

How this metric is used for quality control is shown in Supplementary Fig. 9c. The overlay of ground truth (target) and prediction can act as a direct visual readout of the IoU score. Here, the white areas represent where the signal predicted by the StarDist network overlaps with the target, therefore agreeing with the ground-truth. A large proportion of white areas in the overlay suggests a good agreement between ground truth and prediction which is also reflected by a high IoU score of over 0.8.

While using these metrics as QC for the models cannot prevent errors or artifacts from occurring in the prediction, they can characterize the model's performance. This can be exploited to improve model performance as shown in Supplementary Fig. 11. The figure shows an example of how the IoU score in StarDist can aid in identifying a set of training parameters that can accelerate how quickly the model reaches its top performance. Here, we show how the number of epochs, number of steps, and batch size can affect the IoU score. Additionally, the number of nuclei found in the image also provides a way to assess the model performance compared to the number of nuclei identified from the ground-truth (target) mask.

#### 3.4. Mean average precision (mAP) score

In order to quantify the performance of the YOLOv2 models obtained with ZeroCostDL4Mic, we estimate the average precision (AP) of the model on test datasets, as is commonly done to evaluate object detection networks[21,44]. The AP metric encapsulates both how well the model identifies objects in an image and how accurate its class predictions are.

When making predictions, an object detection model will give three outputs per object: the object's bounding box coordinates (identifying the location of the object within the image), its predicted class (what type of object it is) and the confidence (the probability of the class being accurate as estimated by the model for that particular object). The first step to calculate the AP is to rank all object detections of a class by confidence (highest confidence first). The resulting list is then used to calculate the precision and recall parameters with an increasing number of detections taken into account in the ranked order. The precision and recall metrics are defined as follows:

$$Precision = \frac{True\ Positives}{True\ Positives + False\ Positives}$$

$$Recall = \frac{True\ Positives}{True\ Positives + False\ Negatives}$$

In our case, a true positive detection is defined as a detection with the correct class and where the bounding boxes predicted by the model overlap sufficiently with that of the ground-truth annotation for that object. We defined a sufficient level of overlap when the IoU metric (defined in Section 3.3) between ground-truth and predicted bounding boxes reaches a minimum threshold value. By default, in our notebook, this threshold value is set to 0.3.

For each class, the precision and recall scores are then evaluated by including an increasing number of detections following the ranked order, i.e. from highest to lowest confidence. Thus, the precision of a trained model usually begins with values near 1 and tends to drop as more false positives are encountered in the list of detections (lower confidence). In contrast, the recall value measures the proportion of true positives in all possible positives in the dataset which means that it will tend to increase as the number of true positives increases since more detections are taken into account with each object on the list. When these values of precision and recall estimated in ranking order are plotted against each other, this results in a characteristic p-r (precision-recall) curve[45] (see Supplementary Fig. 9d for examples of p-r curves).

This plot aims to represent the typical trade-off between precision and recall (or rather between false positive rate and false-negative rate), since typically, precision will drop as recall improves. One way to interpret the curve is the following: the higher the p-r curve as the recall increases, the better the performance of the model. A perfect model will display a precision of 1 all the way to a recall value of 1. **Therefore, the area under the curve represents a great way to estimate the overall precision of the model for that particular class**. The AP metric essentially describes the area under the curve of the p-r plot. In our notebook, the AP is calculated as outlined in the PASCAL VOC protocol[46] using a simple interpolation in the precision scores, such that the precision at recall $r$ is equal to the maximum precision for any $r' \geq r$. This reduces the effect of individual detections in the data on the AP[21].

Since recall and precision are values with a range of 0 to 1, the AP value also varies from 0 to 1 where values closer to 1 indicate a better performance of the model on a given object class. The mean average precision (mAP) of a YOLOv2 model takes into account the AP of all object classes as follows:

$$mAP = \frac{1}{n}\sum_{i}^{n} AP_i$$

where $n$ is the number of classes in the dataset and $AP_i$ is the average precision of a specific class $i$.

The p-r curve and the AP score for each class as well as the mAP value are all calculated and presented to the user as output in the YOLOv2 notebook QC section. Examples of p-r curves are shown in Supplementary Fig. 9d for the most common cell shape, *elongated*, as also shown in our example dataset (see also in Supplementary Fig. 13). The p-r curves offer a performance comparison between models trained without and with 8x data augmentation (see Supplementary Note 4 for details about data augmentation). The performance of the model significantly improves with 8x data augmentation, as highlighted in the p-r curve by the higher level of precision across the range of recall and the corresponding AP values compared to the non-augmented dataset. (Supplementary Fig. 9d).

For an additional visualization of the QC, the bounding box coordinates for both ground-truth and predicted labels are also saved in .csv files and can be plotted after the quality control step, e.g. using ImageJ.

**Supplementary Note 4: Data augmentation**

Data augmentation is a strategy used to artificially increase the size of training datasets. It commonly consists of applying a set of spatial transformations to both source and target data in the training dataset, such as rotation or vertical/horizontal flipping but more complex transformations can also be used, such as shearing. For instance, simply flipping (horizontal and vertical) and rotating all the images in a dataset by 90 degrees will increase the size of a dataset by 8. Data augmentation may improve the generalization of a model by amplifying diversity in the dataset. This may be especially useful if the available dataset is small, which can occur if it is expensive or time-consuming to generate. All ZeroCostDL4Mic notebooks contain the possibility to enable or disable data augmentation (Supplementary Fig. 12), but its implementation strategy is adapted to each notebook. For instance, we took advantage of the Augmentor library[47] in the StarDist 2D, pix2pix and CARE 2D notebook, while simpler augmentation strategies such as flipping and rotation are implemented in the CARE 3D and fnet notebooks. In the 3D U-Net and the 3D StarDist notebooks, we also implemented the "elastic deform library" (https://pypi.org/project/elasticdeform/). A different augmentation library (imgaug) is used to augment images and bounding boxes in the YOLOv2 notebook (https://github.com/aleju/imgaug)[48].

Importantly, data augmentation can also be detrimental to the training process and lead to the generation of artifacts. So, we recommend that networks be trained with and without augmentation, and for the user to use the QC section available in our notebooks to assess if using data augmentation leads to performance improvements. For instance, when training YOLOv2, with our test dataset, we found that the model performance considerably improved when performing data augmentation on the training images and bounding boxes by flipping and successive rotations by 90 degrees, as quantified by the mAP, shown in Supplementary Fig. 13c and the respective p-r curves shown in Supplementary Fig. 9d. In practice, YOLOv2 models trained on datasets with increasing augmentation factors become more sensitive to the cells in the image and improve bounding box positioning around detected objects (Supplementary Fig. 13a-b).

### Supplementary Note 5: Transfer learning

The performance and generalization of DL networks often scale with the size and diversity of the training dataset. Because of this, trained models often reach peak performance when generated using large training datasets. Importantly, training models with large training datasets require vast amounts of computational power, making them expensive to produce (which in some cases may not be possible within ZeroCostDL4Mic, see Supplementary Note 6 for details on the resource available with Google Colab). Because of this, training very general models is currently limited to computer science developers with access to large resources. On the other hand, individual scientists may want, instead, to train DL models that are high-performance but specific for their data.

However, when publicly available, DL models trained on large amounts of data can be extremely valuable. On the one hand, such pre-trained models can be directly used by users to perform predictions. This can have several downsides and it is not something we would generally recommend (see Supplementary Note 1 for discussion on the topic). On the other hand, a very efficient way to take advantage of these while retaining good performance for the specific data of interest is to use these pre-trained models as a starting point for training a new DL model.

Indeed, a pre-trained model, when trained on large amounts of related data, will contain useful features that can be reused to speed up the training of another model. Therefore, it can be beneficial to use pre-trained models as a starting point for training a DL network instead of starting from a blank model (training from scratch), where all the model weights are typically initialized to randomly allocated values. So, initializing the model weights to those of a pre-trained model is a powerful approach known as transfer learning[49] and is now common practice in the DL field. Transfer learning can improve performance (necessitating fewer epochs) and generate higher-quality models than those trained from scratch[50].

It is also essential to consider that DL approaches have a meaningful environmental impact and transfer learning may help reduce carbon footprints[51]. These advantages make transfer learning an attractive starting point for new custom tasks.

As we believe that transfer learning is a compelling strategy to generate high-quality DL models, we implemented the possibility to perform transfer learning in all of our ZeroCostDL4Mic notebooks. Indeed, users can quickly load pre-trained model weights and re-training these models on custom data (Supplementary Fig. 14). Pre-trained models of broad interest can even be directly downloaded within our notebooks.

To illustrate the performance improvement that can be achieved using transfer learning, we compared the results obtained by training a StarDist model from scratch to the result obtained when a pre-trained model is used as a starting point. The pre-trained model we choose is the *2D_versatile_fluo* StarDist model made available by the StarDist authors[17,18]. This model was generated using data related (images of nuclei) but distinct from our own and is starting to be widely used by the community, to perform prediction, via its integration to the StarDist Fiji plugin[8,9]. Importantly, using the *2D_versatile_fluo* model to perform prediction on our data led to nuclei segmentation but also to the generation of large artifacts rendering it unusable on its own (Supplementary Fig. 14). Typically, to obtain high-quality predictions from models trained from scratch, we needed to train our StarDist models for more than 200 epochs (Supplementary Fig. 14). However, when using the *2D_versatile_fluo model* as a starting point via transfer learning, very high-quality prediction can be made using a model trained for as little as 5 epochs (Supplementary Fig. 14).

Transfer learning also allows the user to circumvent the Google Colab 12h training time limit (see Supplementary Note 6 for discussion on resources available with Google Colab). Loading a pre-trained model for training also enables training to occur over multiple Colab runtimes, as well as to do transfer learning using models trained outside of ZeroCostDL4Mic. We believe that transfer learning is an especially attractive feature of ZeroCostDL4Mic as it allows users to tune generalist models to their data and generate optimal results while requiring minimal resources and time. Therefore we encourage developers and users to share their trained models via the emerging online “model zoo” so that they can be used by others to enable faster re-training.

**Supplementary Note 6: Capabilities of Google Colab**

The Google Colab (https://colab.research.google.com) platform offers free and easy access to a Graphical Processing Unit (GPU) and Tensor Processing Unit (TPU), which enables users to train many networks on bespoke training datasets, as well as running predictions on unseen data. In practice, for each notebook session, Google Colab assembles a virtual machine with allocated RAM, disk space, and access to GPU/TPU. Although these resources are free, they are finite. It is important to consider the available resources when exploiting Google Colab for DL training and predictions. In our experience, we consistently found that GPU acceleration provided faster computation than TPU for the networks and datasets presented here. Therefore, we focused our attention on GPU accessibility and performance below. In the following sections, we discuss how to handle these resources in order to perform efficient training.

**6.1. Google Drive storage**

When using a free Google Drive (https://www.google.com/drive/) account to perform training, the user will have access to 15 GB of data storage that can be accessed by any Google Colab notebook. All training and test datasets, plus the output of the training, need to fit within this 15 GB limit. We have shown, however, that this is sufficient to efficiently train all of the networks with the datasets that we provide (see Supplementary Table 1). Additional storage space can also be purchased from Google.

**6.2. Remote RAM capacity**

A 12.72 GB RAM limit currently exists for the free GPU or TPU provided by Google Colab. The amount of RAM required to execute the code is determined by the size of the training data, the number and sizes of patches/batches. Exceeding this RAM limit can cause the notebook to crash when initializing the networks. For the datasets we have tested, it was always possible to train the networks with the currently available RAM. It is important to note that large datasets (e.g. datasets made of large 3D stacks) may reach or exceed the RAM capacity when using a large number of patches/batches or data augmentation.

#### 6.3. Time-outs

***12-hour time-out.*** The time taken to train a network sufficiently is primarily determined by the size of the training dataset, the number of steps/epochs/patches, and the efficiency of the underlying code. Google Colab currently offers 12 hours of GPU/TPU access after which remote data loaded into the virtual machine will be lost, a limit primarily enforced to prevent cryptocurrency-mining. For users, this means that if training has not completed by the 12h limit, it will be stopped. This constraint can be annoying when networks need to be trained over many epochs to reach high performance, often necessary for large datasets. However, users can easily circumvent this issue by continuing their training sessions over multiple Colab runtime. Indeed, in all ZeroCostDL4Mic notebooks, model checkpoints are automatically saved, in Google Drive, during training which allows the users to continue training later from such a checkpoint (see Supplementary Note 5 for discussion on transfer learning).

***Log-out if idle.*** Google Colab may disconnect significantly earlier than after 12 hours if it detects an interruption of user interaction or network training. Usually, this time-out happens after 30 to 90 minutes of idleness in our hands, i.e. code not running or lack of user interaction with the Google Colab interface. When the log-out occurs, local variables, installed packages and data stored outside any mounted drive are deleted. Hence, if the log-out occurs before training a model, cells setting up parameters for training such as paths or hyperparameters may have to be reset before training. With all of our notebooks, the models are automatically saved in Google Drive upon training completion, meaning that long training sessions do not have to be attended.

#### 6.4. Inconsistent GPU access

Google Colab does not guarantee access to a GPU, as the number of current users may be larger than the number of available devices. It is not clear how access to a GPU is regulated, but it may be determined by traffic to the Google Colab servers. If no GPUs are available, Colab will offer to run the notebook using either a TPU, CPU (without acceleration), or as a local runtime (i.e. on the machine of the user). It should be noted though that these alternatives can be significantly slower than Google Colab GPU access. However, GPU access usually becomes available again from within a few hours to a day.

#### 6.5. GPU type

Google Colab uses different GPUs that currently include NVIDIA K80, T4, P4, and P100 (as of August 2020). The user cannot decide which GPU will be available when using the notebook. According to the Google Colab (https://research.google.com/colaboratory/faq.html) FAQ, this is due to limitations in the provision of a free service to users which makes certain types of GPU unavailable at the time a notebook is used. In practice, this does not affect the performance of the models trained with ZeroCostDL4Mic. However, it will affect the speed at which networks can be trained and used. Therefore, users might encounter variability in training times as a consequence. To find out which type of GPU Google Colab is using, the user can play the first cell in each notebook which will give information on GPU access and type.

#### 6.6. How to best handle Google Colab resources

Several steps can be taken by users of ZeroCostDL4Mic to optimize the usage of Google Colab's resources. Regarding the 12h maximum training time, we encourage users to change the number of epochs and training parameters so that the training takes less than 12h. This should be possible with all of the networks we provide. Nevertheless, we also provide the option to continue the training from a saved training checkpoint in case of a time-out. This allows the user to start from a pre-trained model and accumulate many rounds of steps (see Supplementary Note 5 about transfer learning).

For the log-out if idle issue, Google Colab cells can be played all at once (or a subset at once). In this case, the activated cells will run one after the other. This can be useful to ensure that the runtime does not disconnect until the completion of all the analytical steps and the user data saved in Google Drive.

ZeroCostDL4Mic is aimed to be an entry point to learn about DL methods where users can quickly train networks with their data. While ZeroCostDL4Mic, associated with the Google Colab platform, is entirely free to use, the resources available can be easily extended with small financial investments. For instance, the free 15 GB Google Drive storage space can be increased by purchasing more Google Drive storage from Google. In addition, Google is rolling out a Google Colab Pro (https://colab.research.google.com/signup#) version that offers faster GPUs, longer runtimes and access to more RAM. If these intermediate options are still not sufficient, ZeroCostDL4Mic notebooks can also be adapted to run on the user's own computer by connecting Google Colab to a local runtime (https://research.google.com/colaboratory/local-runtimes.html). This option allows the user to access their local files and resources (local GPU) directly from the Google Colab notebooks. This, of course, requires the users to invest in a powerful workstation.

#### 6.7. Data privacy and ZeroCostDL4Mic

It is important to note that using ZeroCostDL4Mic may not always be suitable for the analysis of confidential data as the images need to be uploaded to Google Drive prior to analysis. Therefore, we advise users to read the general conditions of using a Google Drive account https://www.google.com/drive/terms-of-service/ before using our notebooks in case concerns about data protection exist. Importantly, Google claims no ownership rights over content stored in Google Drive and the use of Google Colab is not different. In our hands, removal, editing, or organization of files have not occurred when developing and testing ZeroCostDL4Mic. According to Google Colab terms of use, file modifications become more likely if Google's terms are breached (https://support.google.com/docs/answer/148505) or if the user has given specific permission for files to be edited.

To ensure data safety, we recommend our users to upload images without their associated metadata (all the information that details what the images actually are). If this simple precaution is taken, we do not foresee that data privacy issues would affect most microscopy image analysis needs, as images without their associated metadata are virtually worthless. It is important to note that users can also upload only the dataset required to train DL networks and perform the prediction locally to alleviate the issues with storing large amounts of sensitive data in the cloud.

**Supplementary Note 7: Future perspectives for ZeroCostDL4Mic**

Together with the help of the wider research community, we expect to grow the number of networks available in ZeroCostDL4Mic quickly. We also expect ZeroCostDL4Mic to become a standard framework which developers can use to showcase and evolve their networks, adapting them to data analysis tasks optimized for their specific image processing problems. To this end, we will continue developing ZeroCostDL4Mic, and further adapting it to novel DL paradigms. This focus will incorporate the capacity to maintain compatibility with the rapidly evolving DL libraries available today and the ability to export pre-trained models which could be used in either DeepImageJ[2], CSBDeep[3] or other prediction engines.

### Supplementary Tables

| Network | Data Type | # of files | Image Sizes | Image Type | Comments |
|---|---|---|---|---|---|
| U-Net | Training - Images | 28 | 512x512 | EM 8-bit TIFF | ISBI or here |
| U-Net | Training - Masks | 28 | 512x512 | Binary 8-bit TIFF | ISBI or here |
| U-Net | Test - Images | 2 | 512x512 | EM 8-bit TIFF | ISBI or here |
| U-Net | Test - Masks | 2 | 512x512 | Binary 8-bit TIFF | ISBI or here |
| 3D U-Net | Training/Test - Images | 165 | 1024x768 | EM 8-bit TIFF | here |
| 3D U-Net | Training/Test - Masks | 165 | 1024x768 | Binary TIFF | here |
| Stardist (2D) | Training - Images | 45 | 1024x1024 | Fluo 16-bit TIF | This paper, and here |
| Stardist (2D) | Training - Masks | 45 | 1024x1024 | Object-labelled 8-bit TIF | This paper, and here |
| Stardist (2D) | Test - Images | 2 | 1024x1024 | Fluo 16-bit TIF | This paper, and here |
| Stardist (2D) | Test - Masks | 2 | 1024x1024 | Object labelled 8-bit TIF | This paper, and here |
| Stardist (2D) | Test - Stacks | 4 | 1024x1024x86 | Fluo 16-bit TIF | This paper, and here |
| Stardist (3D) | Training - Images | 27 | 128x128x64 | Fluo 16-bit TIF | Martin Weigert et al , 2020 (4) and here |
| Stardist (3D) | Training - Masks | 27 | 128x128x64 | Fluo 16-bit TIF | Martin Weigert et al, 2020 (4) and here |
| Stardist (3D) | Test - Images | 3 | 128x128x64 | Fluo 16-bit TIF | Martin Weigert et al, 2020 (4) and here |
| Stardist (3D) | Test - Images | 3 | 128x128x64 | Fluo 16-bit TIF | Martin Weigert et al, 2020 (4) and here |
| YOLOv2 | Training - Brightfield Images | 30 | 1040x1380 | 8-bit PNG | This paper |
| YOLOv2 | Training - Hand-annotations | 30 | n.a. | xml (Pascal VOC - format) | This paper |
| YOLOv2 | Test - Brightfield Images | 3 | 1040x1380 | 8-bit PNG | This paper |
| YOLOv2 | Test - Hand-annotations | 3 | n.a. | xml (Pascal VOC - format) | This paper |
| N2V (2D) | Training - Images | 1 | 512x512 | Fluo 16-bit TIFF | Stubb et al, 2020 (9), and here |
| N2V (2D) | Test - Images | 22 | 512x512 | Fluo 16-bit TIFF | Stubb et al, 2020 (9), and here |
| N2V (3D) | Training - Images | 1 (Actin) +1 (Fibronectin) | 512x512x13 | Fluo 16-bit TIFF | Kaukonen et al, 2017 (10) (Actin and fibronectin datasets), and here |
| N2V (3D) | Test - Images | 48 (Actin) +48 (Fibronectin) | 512x512x13 | Fluo 16-bit TIFF | Kaukonen et al, 2017 (10) (Actin and fibronectin datasets), and here |
| CARE (2D) | Training – Low SNR images | 22 | 1024x1024 | SIM fluo (MIP from 3D stack) 32-bit TIFF | This paper (Filopodia dataset - Maximum projection), and here |
| CARE (2D) | Training – High SNR images | 22 | 1024x1024 | SIM fluo (MIP from 3D stack) 32-bit TIFF | This paper (Filopodia dataset - Maximum projection), and here |
| CARE (2D) | Test – Low SNR images | 2 | 1024x1024 | SIM fluo (MIP from 3D stack) 32-bit TIFF | This paper (Filopodia dataset - Maximum projection), and here |
| CARE (2D) | Test – High SNR images | 2 | 1024x1024 | SIM fluo (MIP from 3D stack) 32-bit TIFF | This paper (Filopodia dataset - Maximum projection), and here |
| CARE (3D) | Training – Low SNR images | 22 | 1024x1024x33 | SIM fluo 32-bit TIF | This paper (Filopodia dataset 3D - stack), and here |
| CARE (3D) | Training – High SNR images | 22 | 1024x1024x33 | SIM fluo 32-bit TIF | This paper (Filopodia dataset 3D - stack), and here |
| CARE (3D) | Test – Low SNR images | 2 | 1024x1024x33 | SIM fluo 32-bit TIF | This paper (Filopodia dataset 3D - stack), and here |
| CARE (3D) | Test – High SNR images | 2 | 1024x1024x33 | SIM fluo 32-bit TIF | This paper (Filopodia dataset 3D - stack), and here |
| Deep-STORM | Training - Simulated SMLM data | 2 | 64x64 | 16-bits/32-bits TIFF | This paper |
| Deep-STORM | Training - Ground truth localizations | 2 | n.a. | .csv file | This paper |
| Deep-STORM | Example dataset - Experimental SMLM data | 2 | 256x256 | 16-bits/32-bits TIFF | This paper |
| Label-free prediction (fnet) | Training - Brightfield | 92 | 512x512x32 | Bright-field confocal 8-bit TIF | This paper, and here |
| Label-free prediction (fnet) | Training – Fluo (mitochondrial marker) | 92 | 512x512x32 | Fluo confocal 8-bit TIF | This paper, and here |
| Label-free prediction (fnet) | Test - Brightfield | 8 | 512x512x32 | Bright-field confocal 8-bit TIF | This paper, and here |
| Label-free prediction (fnet) | Test – Fluo (mitochondrial marker) | 8 | 512x512x32 | Fluo confocal 8-bit TIF | This paper, and here |
| pix2pix | Training - LifeAct Spinning disk Images | 1748 | 1024x1024 | 8-bit PNG | This paper |
| pix2pix | Training - SiR DNA Spinning Disk Images | 1748 | 1024x1024 | 8-bit PNG | This paper |
| pix2pix | Test - LifeAct Spinning disk Images | 5 | 1024x1024 | 8-bit PNG | This paper |
| pix2pix | Test - SiR DNA Spinning Disk Images | 5 | 1024x1024 | 8-bit PNG | This paper |
| CycleGAN | Training - Spinning Disk Images | 164 | 1280x1280 | 8-bit PNG | This paper |
| CycleGAN | Training - (unpaired) Fluctuation based super resolution Images | 164 | 1280x1280 | 8-bit PNG | This paper |
| CycleGAN | Test - Spinning Disk Images | 4 | 1280x1280 | 8-bit PNG | This paper |
| CycleGAN | Test - Fluctuation based super resolution Images | 4 | 1280x1280 | 8-bit PNG | This paper |

**Supplementary Table 1: Overview of the available datasets used for training the networks**. For all networks relying on supervised learning, the test datasets consist of the last two files generated for training. These were set aside for testing and are not part of the training dataset.

| Network Name | # of epochs | # of steps | Batch size | Image dimensions | # of images | # of patches, patch size, patch height | GPU type | Time for training (using these settings) |
|---|---|---|---|---|---|---|---|---|
| U-Net | 200 | 6 | 4 | 1024x1024 | 28 | 4, 512x512 | Tesla P100-PCIE-16GB | 8 min |
| 3D U-Net | 75 | 396 | 1 | 1024x768 | 165 | 132, 256x256, 16 | Tesla P100-PCIE-16GB | 575 min |
| StarDist (2D) | 200 | 12 | 2 | 1024x1024 | 72 | 1, 1024x1024 . | Tesla K80 | 170 min |
| StarDist (3D) | 400 | 14 | 2 | 128x128x64 | 27 | 1, 128x128x64 | Tesla P4 | 400 min |
| YOLOv2 | 40 | 30 | 8 | 1040x1380 | 30 | 1, 1040x1380 | Tesla P100-PCIE-16GB | 45min |
| N2V (2D) | 100 | 61 | 128 | 512x512 | 22 | 392, 64 | Tesla T4 | 17 min |
| N2V (3D) | 100 | 133 | 128 | 512x512x13 | 48 | 392, 64, 8 | Tesla P100-PCIE-16GB | 240 min |
| CARE (2D) | 50 | 31 | 64 | 1024x1024 | 22 | 100, 80 | Tesla P100-PCIE-16GB | 3 min |
| CARE (3D) | 50 | 62 | 64 | 1024x1024x33 | 22 | 200, 80, 8 | Tesla P4 | 90 min |
| Deep-STORM | 100 | 313 | 16 | 64x64 | 20 (frames) | 500, 26x26 | Tesla P100-PCIE-16GB | 50min |
| Label-free prediction (fnet) | n.a. | 50000 | 4 | 512x512x32 | 92 | 64, 64, 32 | Tesla P4 | 8h20min |
| pix2pix | 200 | n.a. | 1 | 1024x1024 | 1748 | 512x512 | Tesla P100-PCIE-16GB | 9h |
| CycleGAN | 200 | n.a. | 1 | 1280x1280 | 164 | 512x512 | Tesla P100-PCIE-16GB | 8h54min |

**Supplementary Table 2: Hyperparameters used to train the networks, GPU types allocated and corresponding training times**. Some hyperparameters are hard-coded in the networks or calculated from the other parameters. n.a.: not applicable.

### Supplementary Figures

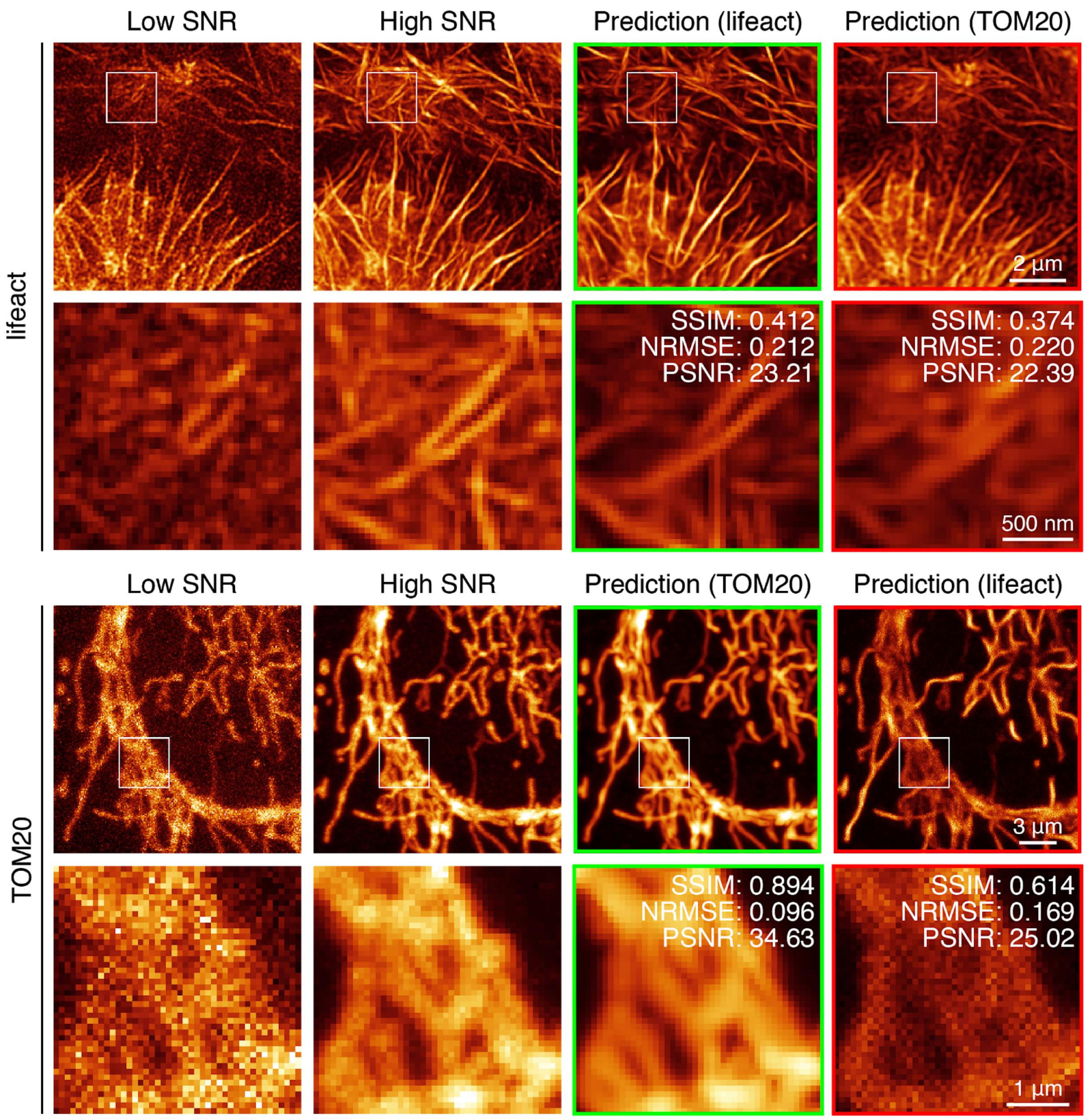


**Supplementary Fig. 1. Of the importance of training models on suitable data. Top:** Filopodia (lifeact) test data and the predictions obtained from a CARE 3D model either trained on similar lifeact dataset (green box) or on TOM20 dataset (red box). **Bottom:** Mitochondria (TOM20) test data and the predictions obtained from a CARE 3D model trained on similar TOM20 dataset (green box) or on lifeact dataset (red box).

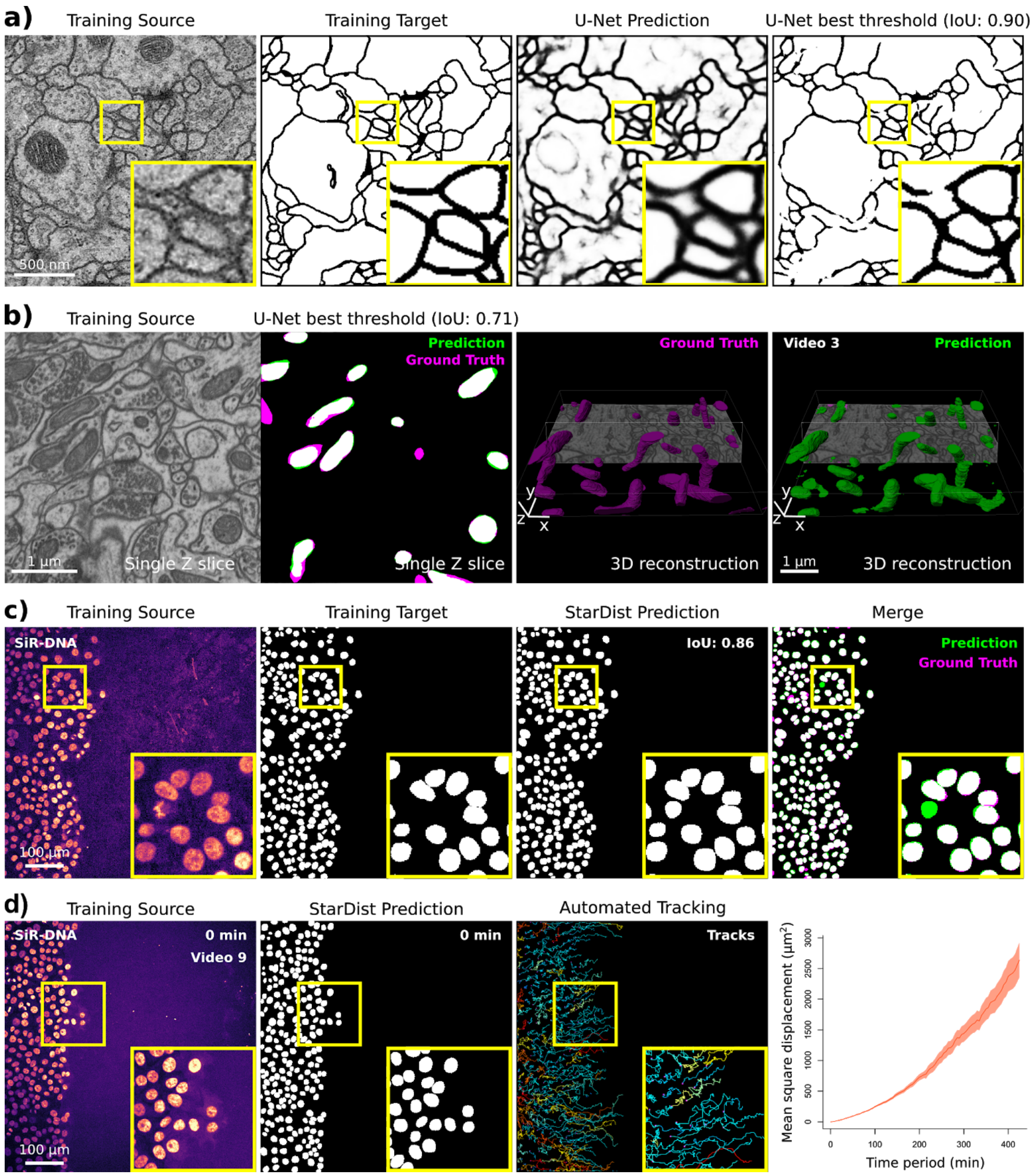


**Supplementary Fig. 2: Image segmentation networks (U-Net and StarDist).** (**a-b**) Example of data that can be generated using the ZeroCostDL4Mic U-Net and StarDist notebooks. **(a)** A 2D U-Net model was trained to segment neuronal membranes from EM images. This training dataset is from the 2012 ISBI segmentation challenge[15]. Training source (raw data), training targets (hand-annotated binary masks), predictions (raw output of the notebook after training and ), and U-Net image thresholded output are displayed. The optimal threshold was assessed automatically using the Quality Control section of the notebook (see Supplementary Note 3). **(b)** A 3D U-Net network was trained to segment mitochondria from EM images. The training dataset was made available by EPFL and consists of EM images of 5x5x5 µm$^3$ sections taken from the CA1 hippocampus region of the brain. A representative single Z slice as well as an overlay displaying U-Net prediction and the ground truth are displayed. 3D reconstructions displayed were performed from U-Net predictions using Imaris (Supplementary Video 8). (**c**, **d**) Example of data that can be generated using the ZeroCostDL4Mic StarDist notebooks. (**c**, **d**) A StarDist model was trained (**c**) to automatically detect nuclei in movies of migrating DCIS.COM cells, labeled with SiR-DNA, to automatically track their movement (**d**). (**c**) Example of Training source (DCIS.COM cells labeled with SiR-DNA), Training targets (Ground truth masks), and StarDist prediction are displayed. (**d**) StarDist outputs were used to automatically track cell movement over time in TrackMate (Supplementary Video 9). Cell tracks were further analyzed using the online platform motilitylab.net.

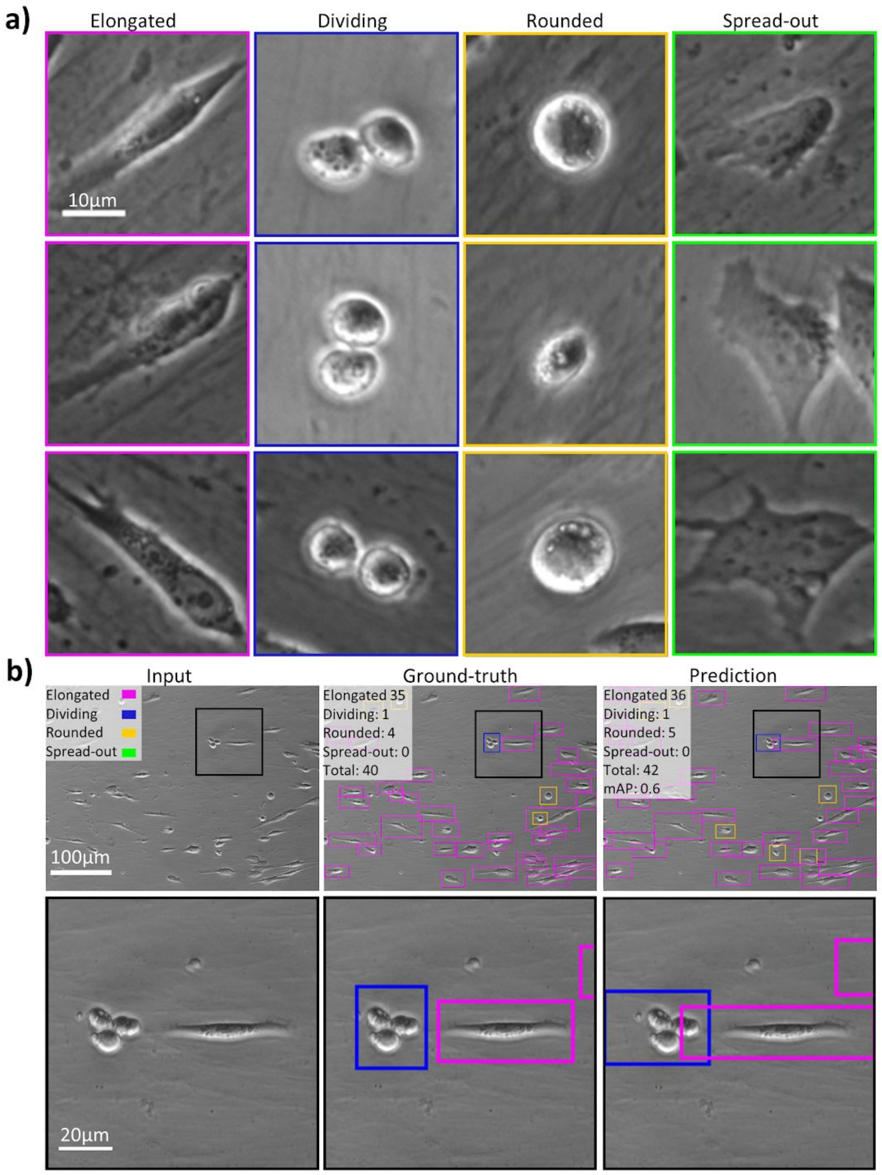


**Supplementary Fig. 3: Object detection (YOLOv2).** Example of data that can be generated using the ZeroCostDL4Mic YOLOv2 notebook, detecting and identifying cell shape classification from a cell migration brightfield time-lapse dataset. **(a)** Identified cell shapes and representative examples that were hand-labeled in the training dataset. **(b)** Input, ground-truth, and prediction obtained from object detection, highlighting the identification of the presence of 3 classes in the field-of-view. mAP: mean average precision (see Supplementary Note 3.4 for details).

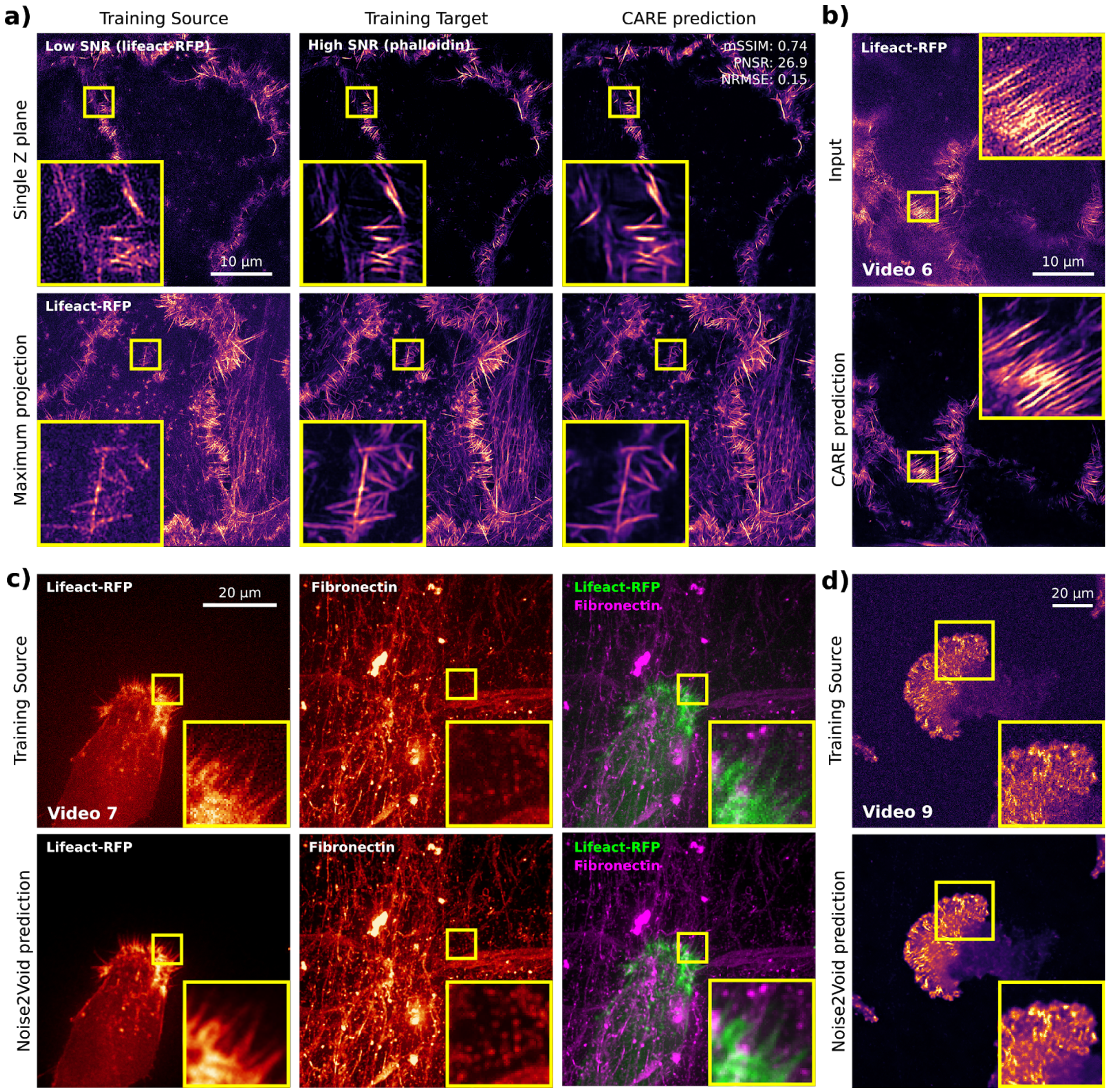


**Supplementary Fig. 4**: **Image denoising and restoration networks (CARE and Noise2Void)**. Example of data that can be generated using the ZeroCostDL4Mic CARE and Noise2Void notebooks. (**a, b**) A 3D CARE network was trained using SIM images of the actin cytoskeleton of DCIS.COM cells using fixed samples (**a**) to denoise live-cell imaging data (**b**). (**c**) Fixed samples were imaged using SIM to obtain low signal-to-noise images (lifeact-RFP, Training Source) and matching high signal to noise (Phalloidin staining, Training Target) images and this paired dataset was used to train CARE. Input, ground truth, and a CARE prediction are displayed (both single z plane and maximal projections). The QC metrics values computed directly in the CARE notebook are indicated. (**b**) The network trained in (**a**) was then used to restore live-cell imaging data (Supplementary Video 6). The low SNR image (input) and the associated CARE predictions are displayed (single plane). (**c**) Movie of an ovarian carcinoma cell labeled with lifeact-RFP migrating on cell-derived matrices (labeled for fibronectin) denoised using Noise2Void. Both training source and Noise2Void predictions are displayed (Supplementary Video 7). For each channel, a single Z stack (timepoint) was used to train noise2Void, and the resulting model was applied to the rest of the movie. (**d**) Movie of a glioma cell endogenously labeled for paxillin-GFP, migrating on 9.6kPa polyacrylamide hydrogel, and imaged using an SDC. Both training source and Noise2Void prediction are displayed (Supplementary Video 9). A single image (time point) was used to train Noise2Void and the resulting model was applied to the rest of the movie. For all panels, yellow squares highlight a region of interest which is magnified.

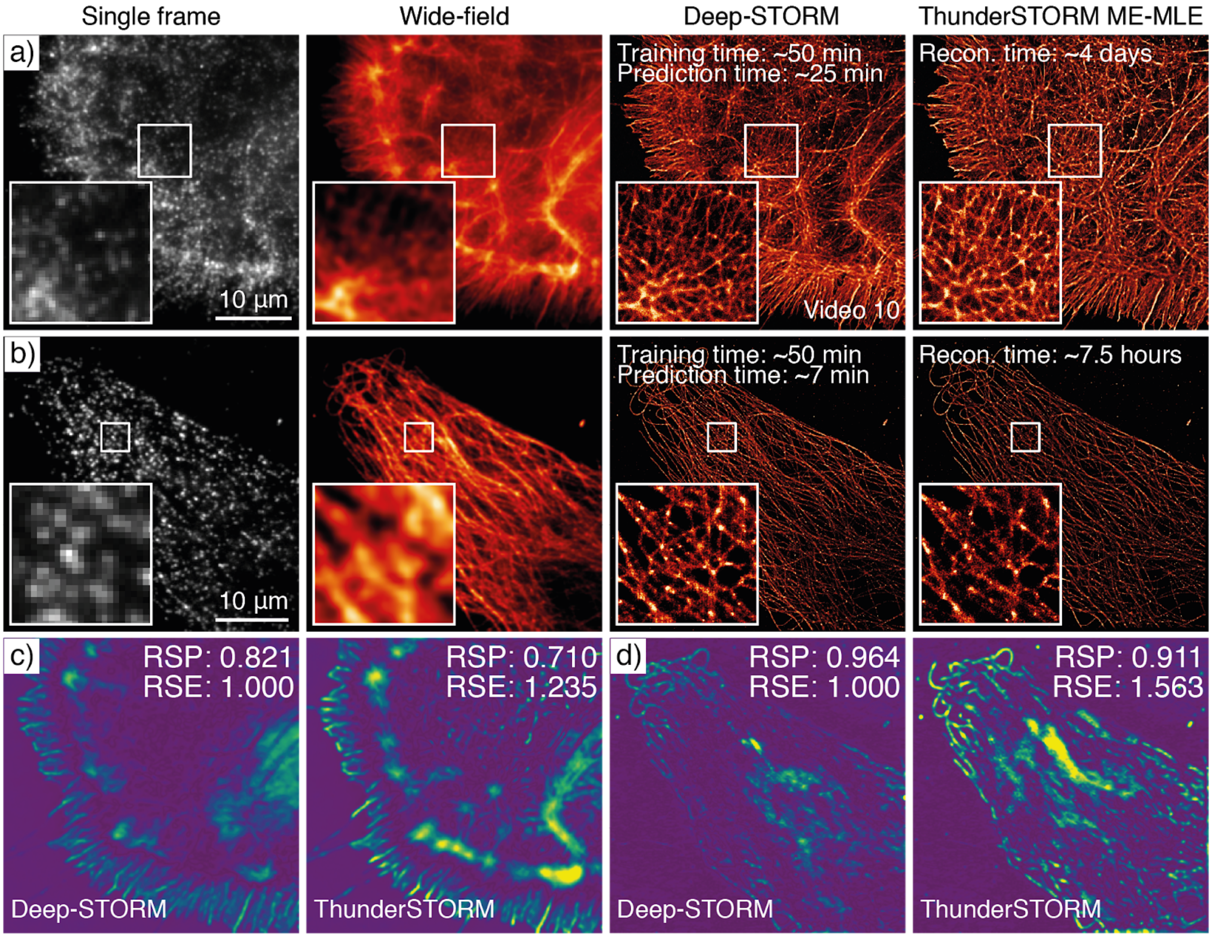


**Supplementary Fig. 5**: **Super-resolution microscopy network (Deep-STORM).** Example of data that can be generated using the ZeroCostDL4Mic Deep-STORM notebook. (a) Deep-STORM reconstruction of the BIN10 dataset, phalloidin labeling of a glial cell, and comparison with ThunderSTORM[34] Multi-Emitter Maximum likelihood estimation (ME-MLE). See also Supplementary Video 10 for more details. (b) Deep-STORM reconstruction of a U2OS cell immuno-labeled for tubulin and imaged via DNA-PAINT[40], as well as the comparison with ThunderSTORM ME-MLE. (c) and (d) SQUIRREL[35] analysis comparing reconstructions of Deep-STORM and ThunderSTORM ME-MLE from a) and b), highlighting a better linearity of the reconstruction with respect to the equivalent wide-field dataset for Deep-STORM. ME-MLE processing times were estimated from an Intel Core i7-8700 CPU @ 3.2GHz, 64GB RAM machine for the actin dataset (panel a), and Intel Core i7-8700 CPU @ 3.2 GHz, 32 GB RAM for tubulin data (panel b). The reconstruction times shown for Deep-STORM were obtained with the NVIDIA Tesla P100 PCIe 16 GB RAM available on Google Colab.

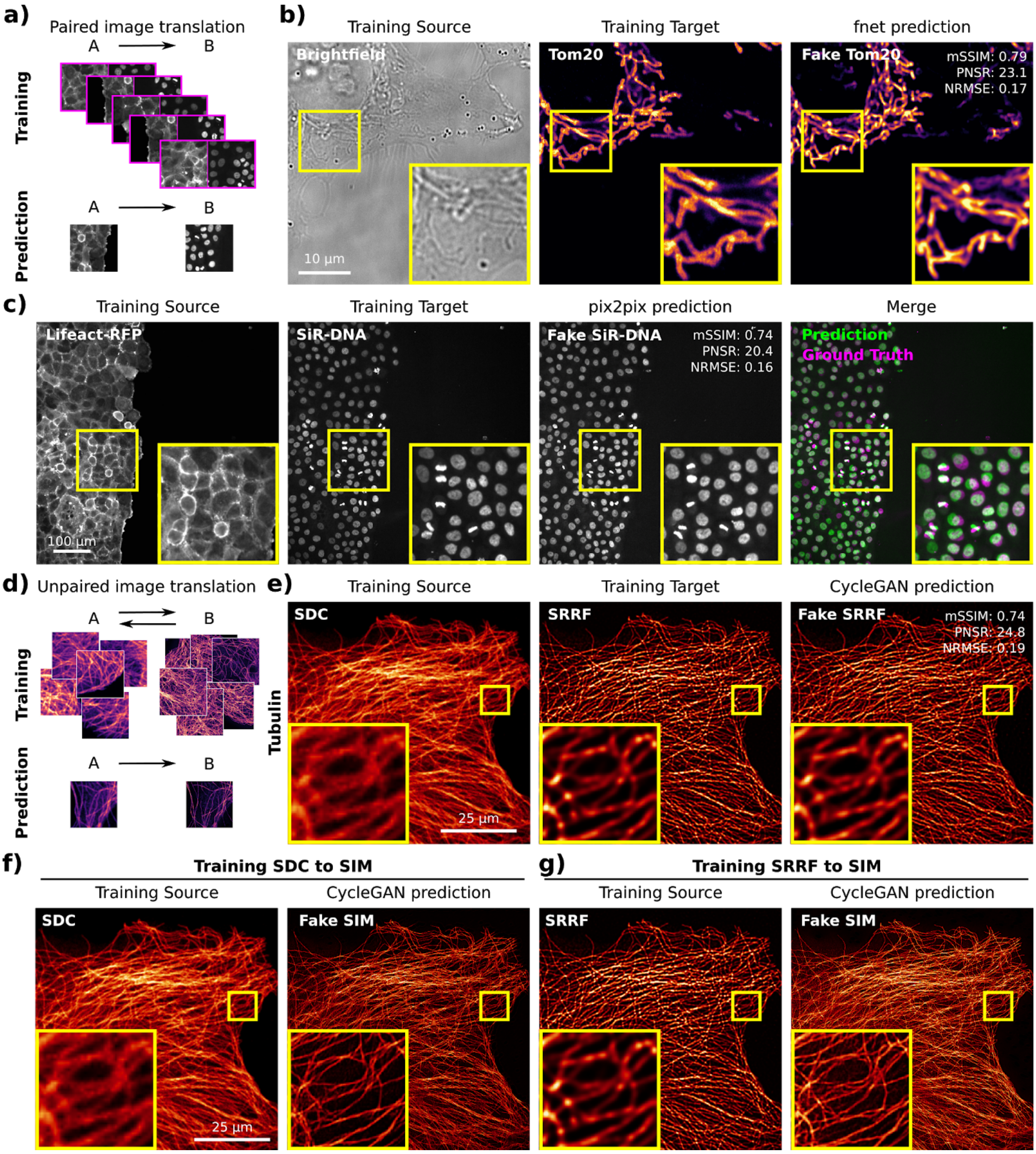


**Supplementary Fig. 6: Image-to-image translation networks (fnet, pix2pix and CycleGAN).** Example of data that can be generated using the ZeroCostDL4Mic fnet, pix2pix and CycleGAN notebooks. (**a**) Scheme illustrating the data required to train paired image-to-image translation networks (pix2pix and fnet). (**b**) Fnet was trained to predict the location of mitochondria (Tom20 staining, Training Target) from brightfield images (Training Source). Both the fnet prediction and the ground truth images are displayed. The QC metrics values computed directly in the fnet notebook are displayed. (**c**) pix2pix was trained to predict nuclear stainings (SiR-DNA, training Target) from actin stainings (lifeact-RFP, Training Source) in migrating DCIS.COM cells. A pix2pix prediction, the corresponding ground truth images are displayed. The quality control metrics values computed directly in the pix2pix notebook are displayed. (**d**) Scheme illustrating the data requirement to train unpaired image-to-image translation networks (CycleGAN). Importantly, these networks do not need to have access to a paired training dataset. (**e, f**) CycleGAN was trained to predict what images of microtubules acquired with a spinning disk confocal (SDC) would look like when processed with SRRF (**e**) or imaged with a SIM microscope (**f**). A CycleGAN model was also trained to transform SRRF images into SIM images (**g**). For the SDC to SRRF translation, the CycleGAN prediction and ground truth SRRF images are displayed as well as the QC metrics values computed directly in the pix2pix notebook are displayed. For all panels, yellow squares highlight a region of interest which is magnified.

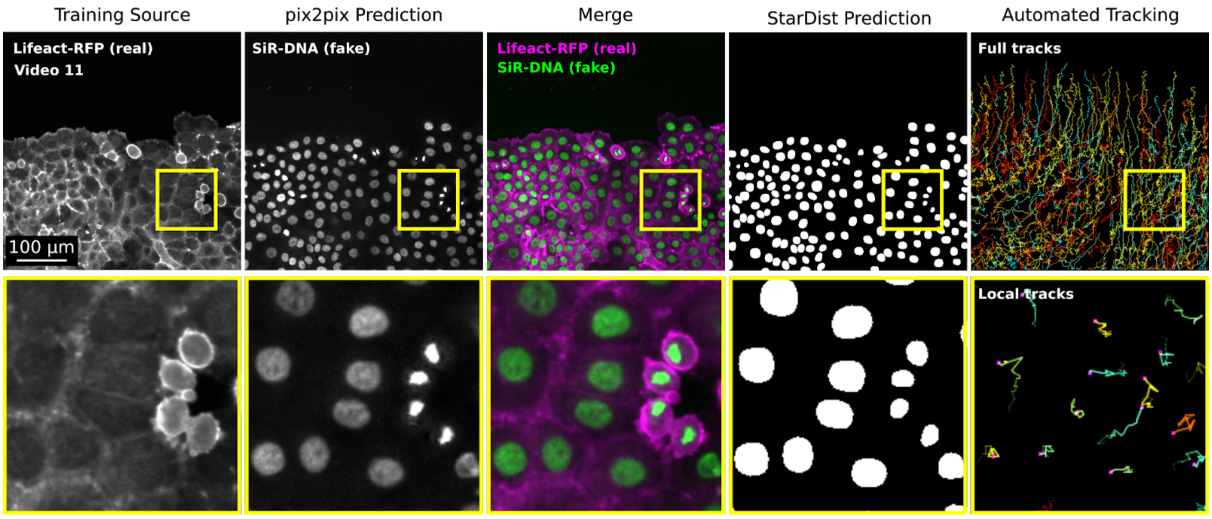


**Supplementary Fig. 7: Example illustrating how ZeroCostDL4Mic notebook can be used together.** Figure highlighting how ZeroCostDL4Mic notebooks can be combined to create a data analysis pipeline. Here we wanted to automatically track the migration pattern of DCIS.COM cells labeled with lifeact-RFP. Therefore we first used pix2pix to predict the actin staining into nuclei staining (as in Supplementary Fig. 6c) and StarDist to detect the nuclei. From the StarDist prediction, cells were automatically tracked using TrackMate[1] (as in Supplementary Fig. 2d; see also Supplementary Video 11). A representative field-of-view is displayed.

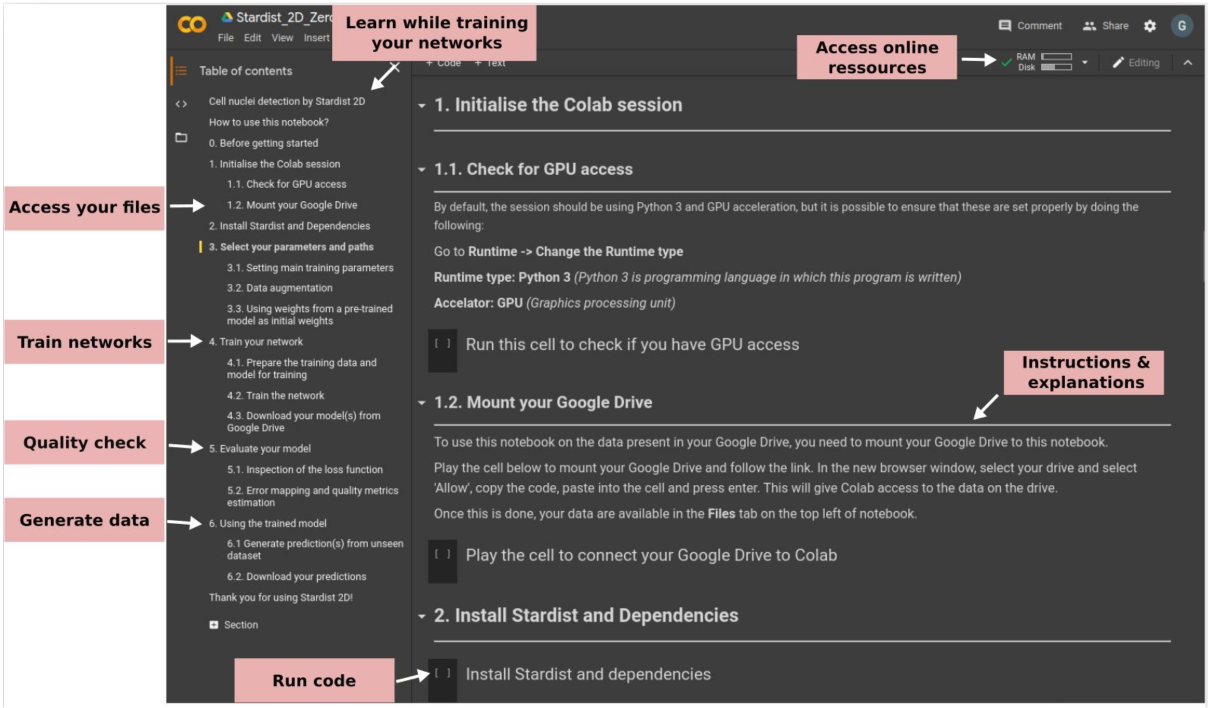


**Supplementary Fig. 8: Graphical user interface (GUI) of the ZeroCostDL4Mic notebooks.** The layout of the notebook and quick access to the different sections is available on the left panel. The user has access to the files present on their Google Drive.

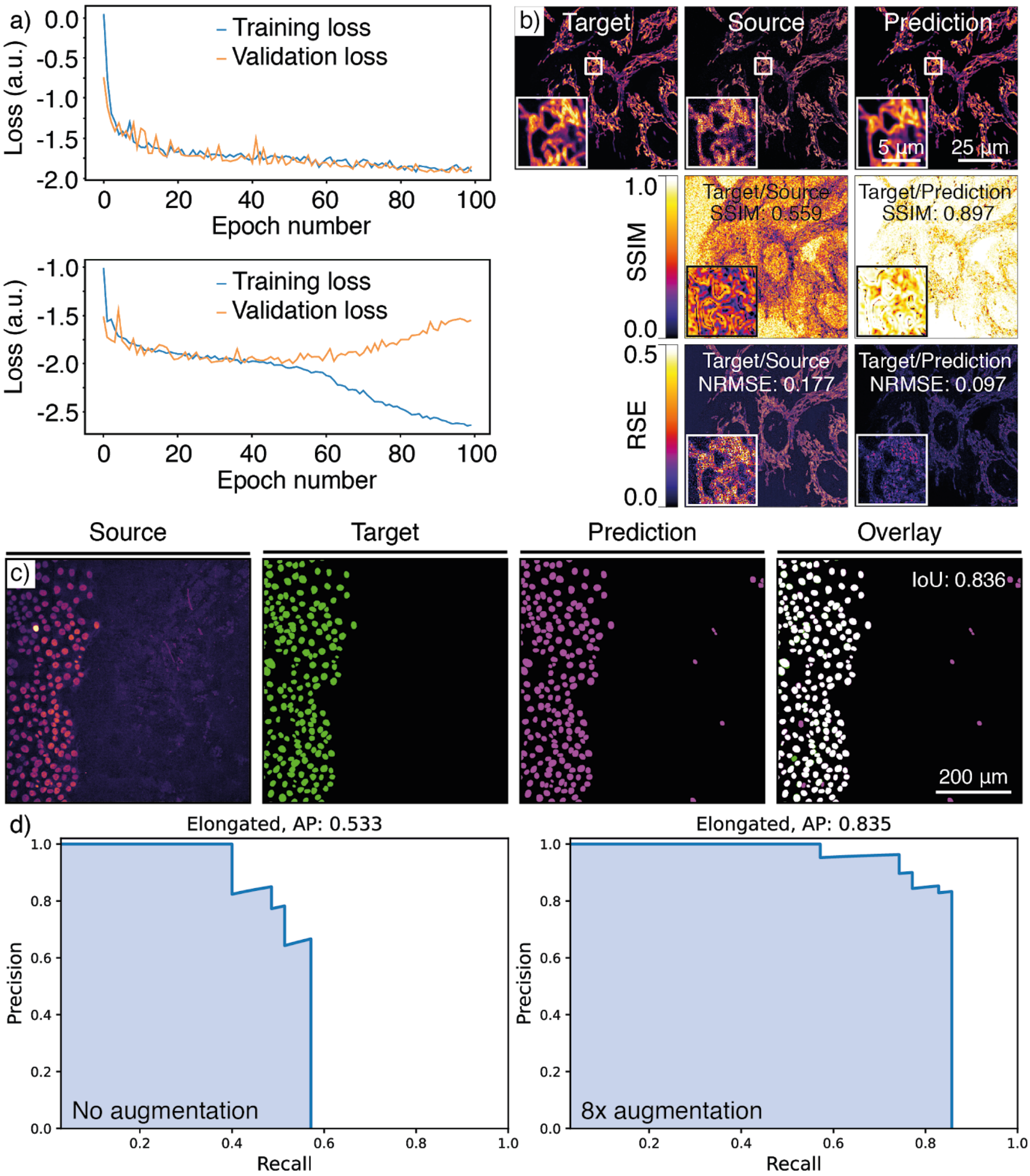


**Supplementary Fig. 9: Quality Control of trained models. a) Overfitting models:** Graphs showing training loss and validation loss curves of a StarDist network with different hyperparameters. The upper panel shows a good fit of the model to unseen (validation) data, the lower panel shows an example of a model that overfits the training dataset. **b) RSE and SSIM maps:** An example of quality control for CARE denoising model performance. **c) IoU maps:** An example of quality control metrics for a StarDist segmentation result. **d) Precision-recall (p-r) curves**: p-r curves for the dataset shown in Supplementary Fig. 13, highlighting the effect of augmentation on the performance metrics of the YOLOv2 model.

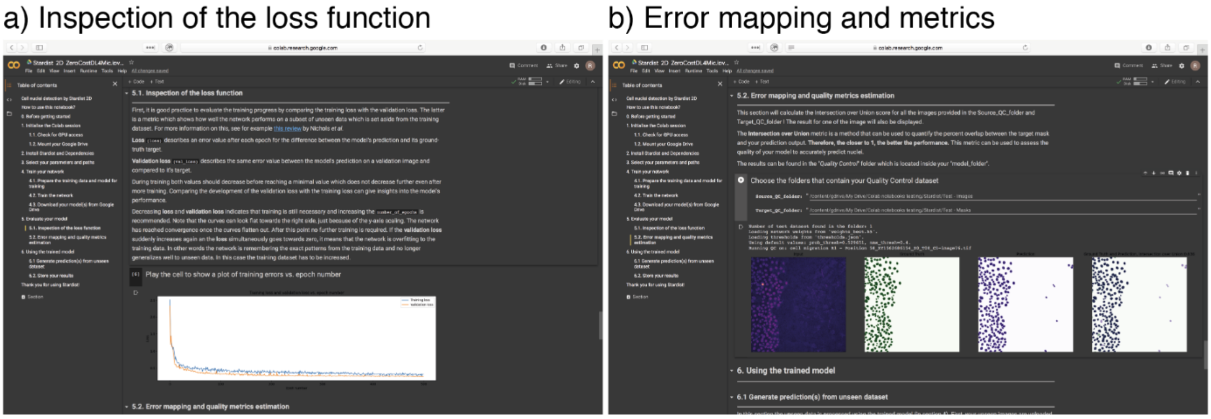


**Supplementary Fig. 10: Quality Control in the ZeroCostDL4Mic notebooks.** Screenshot of the two quality control steps performed in the StarDist notebook. These quality control sections are available in all of the notebooks that we provide.

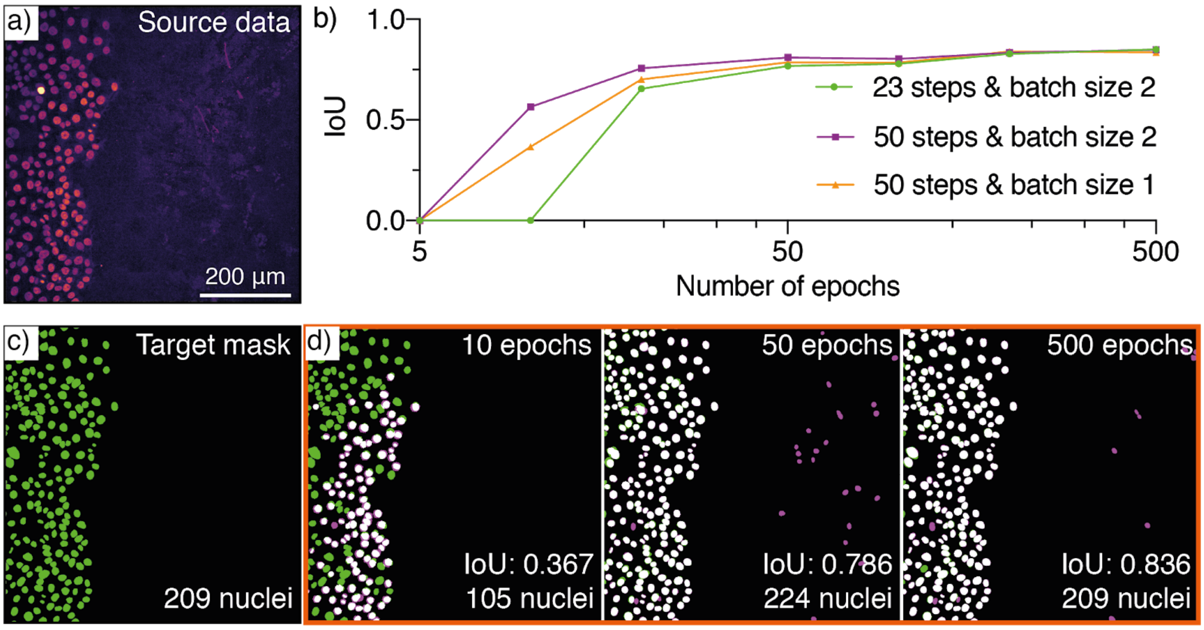


**Supplementary Fig. 11. Model performance vs. training parameters.** This figure displays an example of how adjusting parameters (batch size, number of steps and number of training epochs) can affect model performance in StarDist nuclear segmentation. It is also a demonstration of how assessing trained models in the ZeroCostDL4Mic notebooks can inform hyperparameter choices and improve predictions.

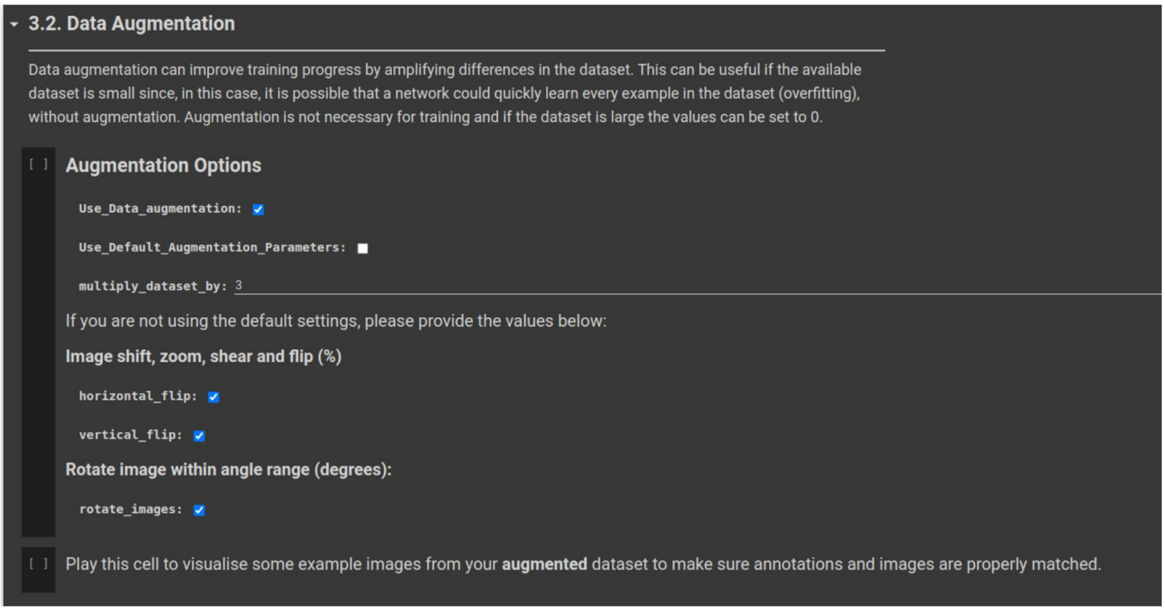


**Supplementary Fig. 12: Example of data augmentation section in the ZeroCostDL4Mic fnet notebook.** Screenshot highlighting the data augmentation section available in the fnet notebook. Here, only horizontal flip, vertical flip and 90-degree rotations are implemented. Data augmentation can be enabled or disabled in all the provided notebooks.

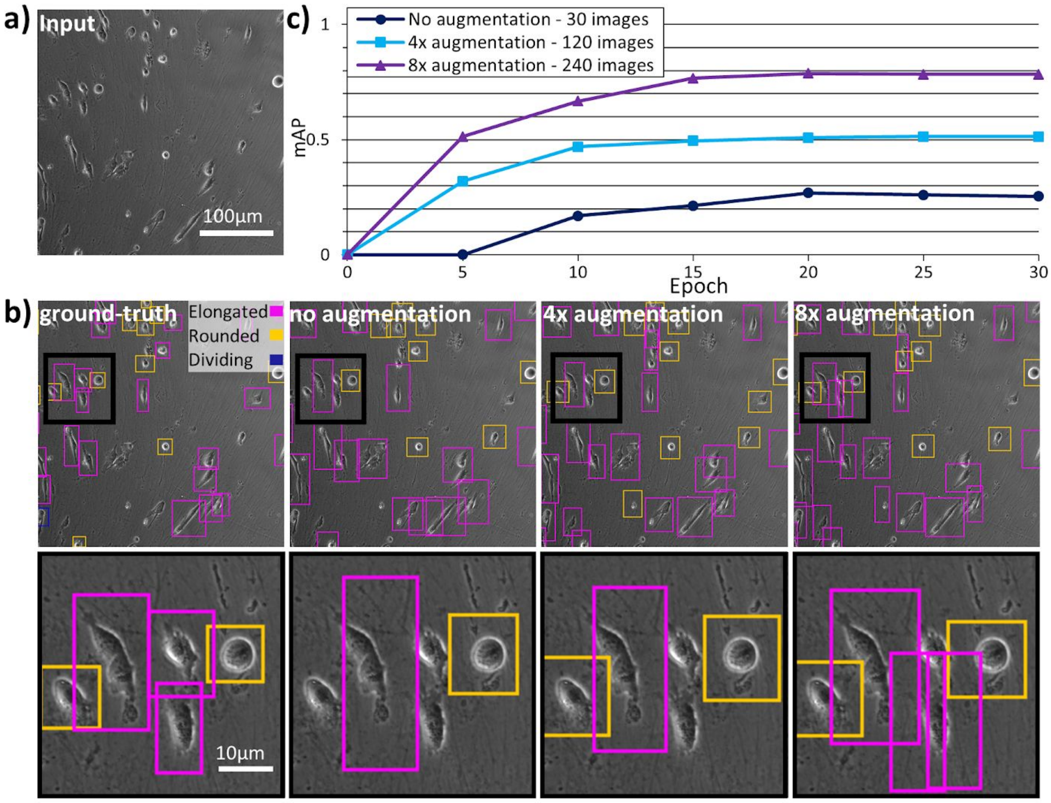


**Supplementary Fig. 13: Data augmentation can improve prediction performance.** YOLOv2 cell shape detection applied to brightfield time-lapse dataset. **(a)** Raw brightfield input image. **(b)** Ground-truth and YOLOv2 model predictions (after 30 epochs) with increasing amounts of data augmentation. The original dataset contained 30 images which were first augmented by vertical and horizontal mirroring and then by 90 degrees rotations. **(c)** Mean average precision (mAP) as a function of epoch number for different levels of data augmentation.

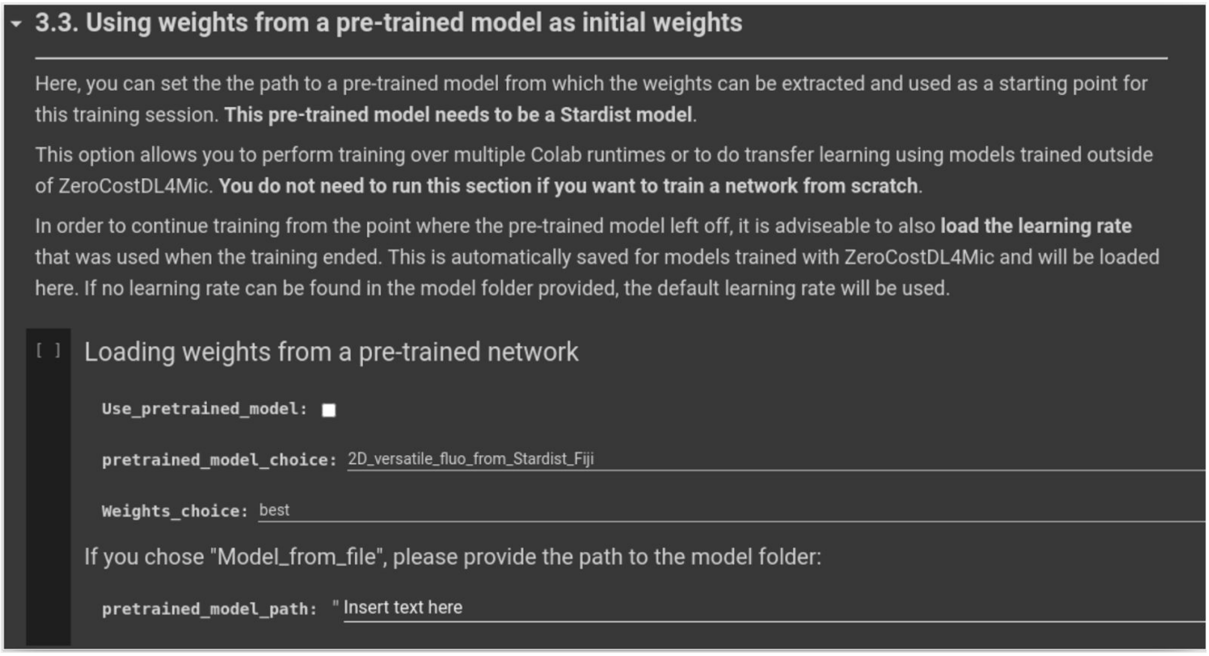


**Supplementary Fig. 14: Transfer learning in the ZeroCostDL4Mic notebooks.** Screenshot highlighting the Transfer learning section available in the CARE 2D notebook. A similar Transfer learning section is provided in all the ZeroCostDL4Mic notebooks.

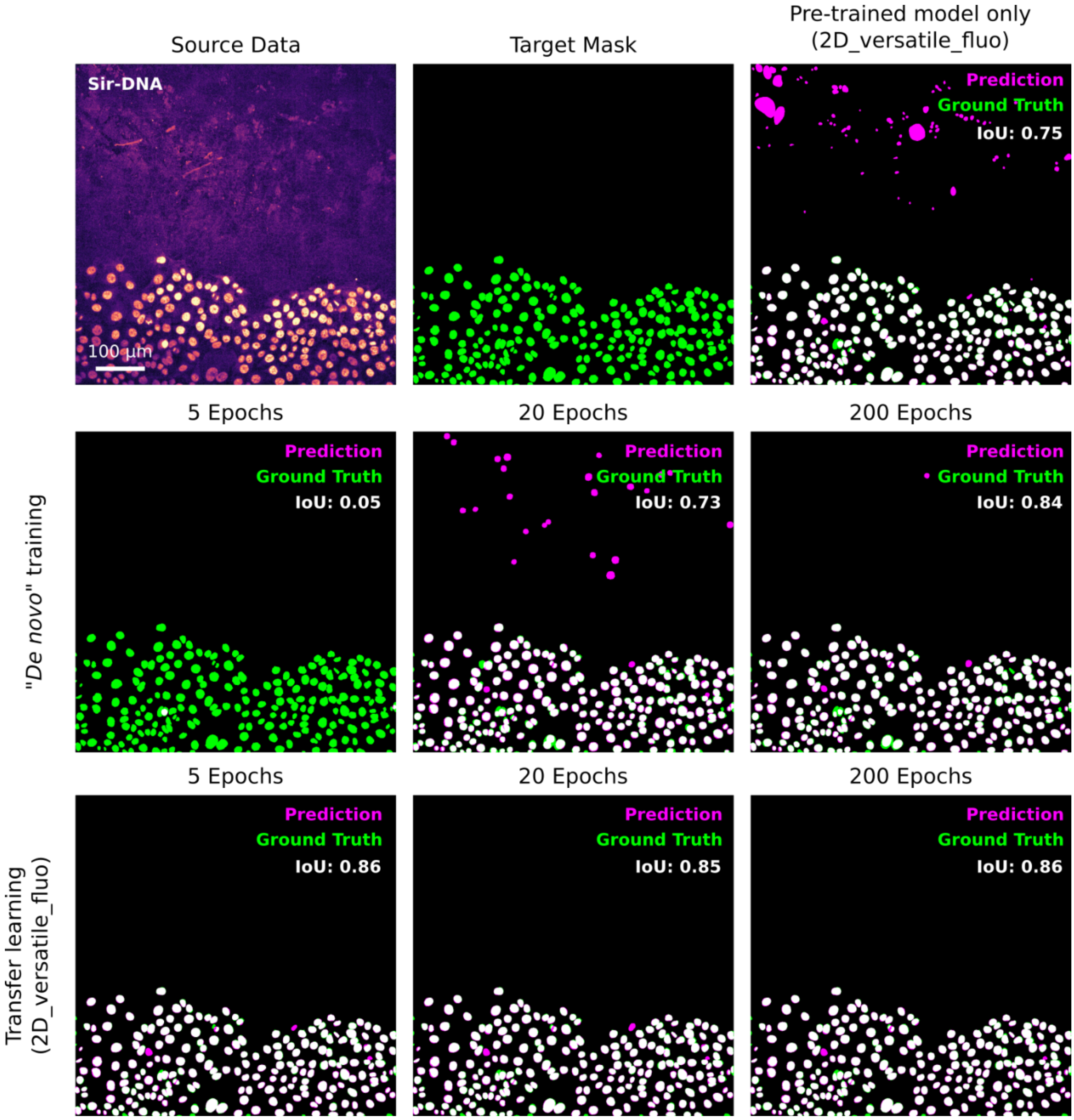


**Supplementary Fig. 15: Transfer learning can reduce training time and improve performance**: This figure displays an example of how transfer learning using a pre-trained model can lead to very high-quality StarDist prediction even after a very short training session. This figure also highlights that using a pre-trained model, even when trained on a large dataset, can lead to inappropriate results.
